# Astrocytic thrombospondins 1 and 2 are required for cortical synapse development controlling instrumental performance

**DOI:** 10.1101/2024.03.01.582935

**Authors:** Oluwadamilola O. Lawal, Francesco Paolo Ulloa Severino, Shiyi Wang, Dhanesh Sivadasan Bindu, Kristina Sakers, Sarah Anne Johnson, Henry H. Yin, Cagla Eroglu

**Author notes:** Corresponding Authors (CE), (HHY).

## Abstract

During development, controlled synaptogenesis is required to form functioning neural circuits that underlie cognition and behavior. Astrocytes, a major glial-cell type in the central nervous system (CNS), promote synapse formation by secreting synaptogenic proteins. Thrombospondins 1 and 2 (TSP1/2), which act through their neuronal receptor α2δ-1, are required for proper intracortical excitatory synaptogenesis. In the adult brain, the loss of α2δ-1 impairs training-induced excitatory synaptogenesis in the anterior cingulate cortex (ACC), and this impairment leads to increased effort-exertion during high-effort tasks. Here, we tested whether TSP1 and TSP2 are required for controlling effort during operant conditioning by using a lever press for food reward training in mice. Surprisingly, we found that constitutive loss of TSP1/2 significantly reduced lever pressing performance when the effort required for a food reward was increased, a phenotype opposite of α2δ-1 loss. Loss of TSP1/2 reduced excitatory synapse number significantly in adult brains. However, in the ACC of TSP1/2 knockout mice, there was still training-induced excitatory synaptogenesis, likely through the upregulation of TSP4, a TSP isoform that is also synaptogenic. Unexpectedly, we also found a significant increase in inhibitory synapse number and function in the ACC of TSP1/2 knockout mice, which was eliminated after training. Finally, we found that astrocyte-specific ablation of TSP1/2 in developing but not adult astrocytes is sufficient to reduce performance during high-effort tasks. Taken together, our study highlights the importance of developmental astrocyte-derived synaptogenic cues TSP1 and 2 in establishing excitatory and inhibitory circuits that control effort during operant conditioning in adults.

## Introduction

Synapses are the sites of neuron-to-neuron communication in the central nervous system (CNS). Synapse formation and plasticity are critical for the proper development of neural networks that underlie cognition, learning, and memory [1–5]. Synaptic dysfunctions are linked to disorders associated with altered cognition, such as autism spectrum disorder, obsessive-compulsive disorder, and schizophrenia [6–13]. Although synapses are formed between neurons, astrocytes, a glial cell type, play critical roles in developing and remodeling synapses [14–16].

Astrocytes are morphologically complex cells that perform numerous functions in the CNS, including maintaining the blood-brain barrier, providing metabolic support to neurons, and regulating extracellular ion concentrations [17–21]. Though previously they were thought to provide only support to neurons, research in the past few decades has shown that astrocytes regulate synapse formation, function, plasticity, and behavior [16,22–25]. In particular, astrocytes induce synaptogenesis through secreted proteins and cell adhesion-based cues [26–42]. However, how astrocytes control behavior through the secretion of synaptogenic factors is still not well understood.

Thrombospondins (TSPs), a family of oligomeric extracellular matrix proteins, induce excitatory synapse formation between neurons in culture [29,43]. There are five isoforms of thrombospondins (TSP1 – TSP5) [44], and TSP1-4 are expressed in the brain by astrocytes [45–47]. TSP1, TSP2, and TSP4 promote neurite outgrowth [48–50]. TSPs 1 and 2 (TSP1/2) are expressed in the cortex, superior colliculus, and retina during early development in the mouse brain. However, their expression is diminished by postnatal day 21 (P21).

TSPs were discovered as astrocyte-secreted synaptogenic proteins during an investigation to identify the components of astrocyte-conditioned media (ACM) that promote synapse formation [29,43]. In neuron-only cultures addition of TSPs increase structural synapses that are presynaptically active (i.e. releases glutatmate) but are postsynaptically silent due to a lack of 2-amino-3-(5-methyl-3-oxo-1,2-oxazol-4-yl) propanoic acid (AMPA) receptors at the synapse [29]. Previously, it was shown that mice lacking TSP1 and TSP2 (TSP1/2-KO) have significantly fewer cortical excitatory synapses by the third week of postnatal development (P21) [29].

Astrocyte-secreted TSPs induce synaptogenesis by binding to their shared neuronal receptor, α2δ-1, a subunit of voltage-gated calcium channels widely expressed in the nervous system [43]. TSPs possess epidermal growth factor (EGF)-like domains that bind to the Von Willebrand Factor A (VWF-A) domain of α2δ-1 at the post-synapse to activate Rac1 and promote synapse formation and spine maturation [43,51]. Mice lacking α2δ-1 have a significantly reduced number of excitatory synapses and mature dendritic spines in the visual cortex at postnatal day 21 and 40 [51]. Importantly, an anti-epilepic drug called gabapentin, which specifically binds to α2δ-1, disrupts TSP-induced excitatory synapse formation [43,51]. However, whether TSPs induce synaptogenesis in the adult brain and contribute to adult cognitive functions is largely unknown.

Under normal conditions, adult astrocytes are not predicted to express TSP1, and TSP2. However, emerging evidence suggests that their expression may be upregulated in the adult brain and contribute to behavior. For example, in the adult mouse cortex, TSP1/2 expression increases after ischemic stroke [52]. TSP1/2-KO mice have impaired synaptic recovery after stroke, suggesting that TSP1/2 may be important for synaptic plasticity post-injury in the adult brain [52]. Furthermore, chemogenetic activation of Gi protein-coupled receptors in striatal astrocytes increases *Thbs1* mRNA expression and causes hyperactivity with disrupted attention [53]. Blocking the TSP receptor α2δ-1 with gabapentin was able to reverse the behavior phenotype, suggesting that TSP1 upregulation due to astrocytic G protein signaling may underlie behavioral phenotypes [53]. However, it is unknown whether TSP1 or its close homolog TSP2 are required for behaviors.

Synapse formation still occurs in the adult brain and is critical for learning and performance of voluntary behaviors [54–58]. Previously, we investigated whether synaptogenesis occurs during instrumental operant training (lever press for food reward task). During this task, mice first learn to associate that a single lever press results in a food pellet reward. Once the association is formed, we can manipulate the relationship between the lever press and the food reward to understand the mechanisms that underlie different aspects of instrumental learning. For example, we can increase the number of lever presses required for a single food pellet and quantify the mice’s lever press rate and patterns to determine the amount of effort the mice are willing to spend to achieve the reward [59–62]

Previously, we identified a specific region of the prefrontal cortex (PFC), the anterior cingulate cortex (ACC), that was activated during performance of this instrumental task. This activation was found to be due to an increase in the number and activity of excitatory synapses in the trained compared to untrained mice [54]. The loss of the thrombospondin receptor, α2δ-1, ablated training-induced excitatory synapse formation in the ACC but did not impair the learning of the operant task [54]. Instead, α2δ-1 KO mice exhibited a significant increase in effort exertion during high-effort tasks, revealing that they were less sensitive to behavioral cost. Ablation of α2δ-1 specifically in the adult ACC neurons projecting to the dorsomedial striatum, a critical circuit for goal-directed actions (GDAs), was sufficient to impair training-induced synapse formation and increase effort exertion [54].

Since α2δ-1 is required for adult synaptogenesis to control effort exertion, we wondered whether its astrocyte-secreted ligands, TSP1 and TSP2, are also involved in this process. Thus, we investigated whether TSP1/2-KO mice have impaired training-induced synaptogenesis and increased effort exertion, as observed in animals with global deletion of TSP receptor α2δ-1. Unexpectedly, the lack of TSP1/2 reduced performance during high-effort tasks, a phenotype that is the opposite of α2δ-1 KO mice. This result prompted us to investigate the roles of TSP1 and TSP2 in controlling training-induced synapse formation.

## Results

### Loss of TSP1/2 reduces excitatory synapse numbers in the adult ACC

The loss of TSP1/2 [63] was previously shown to impair excitatory synapse formation in the developing mouse cortex [29]. However, whether this deficit persists in adulthood, particularly in the ACC (Fig 1A), is unknown. To determine whether the lack of TSP1/2 reduces excitatory synapse density in layers 1, 2/3, and 5 of the adult ACC (Fig 1B), we utilized a quantitative assay to label and count intracortical excitatory synaptic structures [64,65]. To do so, we quantified the close apposition of pre-synaptic marker, VGlut1, and post-synaptic marker, PSD95, in the adult WT and TSP1/2-KO mice (Fig 1C). This analysis revealed that TSP1/2-KO mice have significantly decreased excitatory synapse density in the adult ACC layers 1, 2/3, and 5 compared to age-matched WT mice of the identical background (Fig 1D-1F).

**Fig 1.**
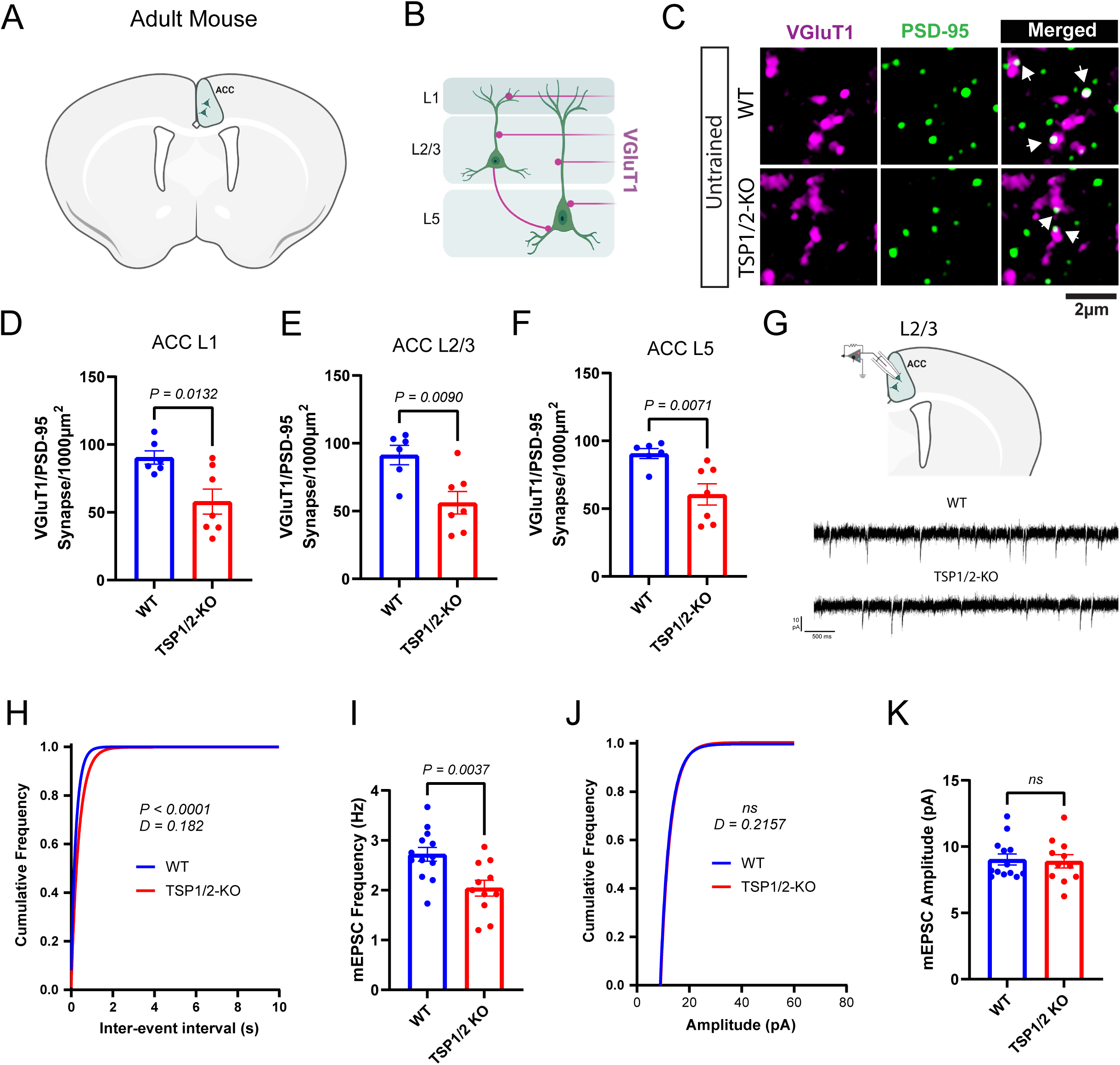
Global loss of TSP1/2 impairs excitatory synaptogenesis in the anterior cingulate cortex. **(A)** Schematic representation of a coronal section of a mouse brain. **(B)** Schematic representation of cortical Layers 1, 2/3, and 5. **(C)** Representative images of VGluT1/PSD-95 staining in the ACC of untrained WT and TSP1/2-KO mice. **(D)** Quantification of VGlut1/PSD-95 co-localized puncta in ACC L1 of untrained mice (n = 6-7 mice per condition; 3 images per mouse). Unpaired Two-tailed t-test. WT (90.46 ± 4.91), TSP1/2-KO (57.94 ± 9.25) [t (11) = 2.95, p = 0.0132]. **(E)** VGlut1/PSD-95 co-localized puncta in ACC L2/3. Unpaired Two-tailed t-test. WT (91.3 ± 7.09), TSP1/2-KO (56.11 ± 8.29) [t (11) = 3.164, p = 0.0090] **(F)** VGlut1/PSD-95 co-localized puncta in ACC L5. Unpaired Two-tailed t-test. WT (90.62 ± 3.66), TSP1/2-KO (60.47 ± 7.83) [t (11) = 3.296, p = 0.0071]. **(G)** Top: Schematic representation of the electrophysiological recordings from ACC L2/3 neurons. Bottom: Example traces from mEPSC recordings from untrained WT and TSP1/2-KO mice. **(H)** Cumulative distribution of the inter-event interval of mEPSC in WT (n = 4 mice; 13 cells) and TSP1/2-KO (n = 4 mice; 11 cells) mice. Kolmogorov-Smirnov test [D = 0.182, p < 0.0001]. **(I)** Average frequency of mEPSC in WT (n = 4 mice, 13 cells; 2.72 ± 0.14 Hz) and TSP1/2-KO (n = 4 mice, 11 cells; 2.04 ± 0.16 Hz) mice. Unpaired t-test [t (22) = 3.25]. **(J)** Cumulative distribution of the amplitude of mEPSC in pA for both WT (n = 4 mice; 13 cells) and TSP1/2-KO (n = 4 mice; 11 cells) mice. Kolmogorov-Smirnov test [D = 0.2157, p = 0.186]. **(K)** Average amplitude of mEPSC in WT (n = 4 mice, 13 cells; 9.03 ± 0.41 pA) and TSP1/2-KO (n = 4 mice, 11 cells; 8.89 ± 0.49 pA) mice. Unpaired t-test [t (22) = 0.222, p = 0.826]. Data shown as mean ± s.e.m.

To determine whether the decreased structural excitatory synapses we observed in the TSP1/2-KOs also cause changes in synaptic function, we performed whole-cell patch-clamp recordings of miniature excitatory postsynaptic currents (mEPSC) from neurons in layers 2/3 of the ACC (Fig 1G). In line with a decrease in synapse numbers, we found a significant decrease in the frequency of mEPSCs (Fig 1H and 1I), but the amplitude did not differ between genotypes (Fig 1J and 1K). Taken together these results show that the loss of TSP1/2 causes a significant decrease in the number and activity of excitatory synapses within the ACC.

This reduction of synapses in the TSP1/2-KO mice could be due to alterations in the cellular composition of the ACC. Therefore, we quantified the number of overall cells (DAPI), astrocytes (Sox9), neurons (NeuN), and oligodendrocytes (Olig2) in the ACC of adult TSP1/2-KO and WT mice (S1A Fig). The loss of TSP1/2 did not change the number of cells in the ACC (S1B-1E Fig). These findings show that TSP1/2-KOs have persisting excitatory synaptic deficits in adulthood in the ACC, a region that regulates learning and performance of goal-directed actions [54,66–71]. Therefore, we next tested whether TSP1/2-KO mice can learn and perform goal-directed behaviors using the lever-press-for-food instrumental operant task.

### Loss of TSP1/2 reduces instrumental performance during high-effort tasks

Previously, we found that TSP receptor α2δ-1 is required in the ACC for excitatory synapse formation that is induced by operant training [54]. These newly formed synapses in the adult ACC are not required for learning but for the regulation of effort exertion, because α2δ-1 global and conditional KO mice pressed the lever at much higher rates for the same reward when compared to WT mice. Since the neuronal receptor for TSP1/2 is α2δ-1, we expected TSP1/2-KO mice will also display the same phenotypes during the instrumental task.

To test this, we trained WT and TSP1/2-KO adult mice in an operant chamber (Fig 2A) to perform the instrumental task on fixed-ratio schedules 1, 5, and 10 (Fig 2B). During fixed ratio 1 (FR1), mice learn to perform one lever press to receive a single food reward. There were no differences in the performance of mice from both genotypes as measured by the number of lever presses per minute (Fig 2C). Surprisingly, however, during FR5 and FR10, when mice must press the lever 5 or 10 times to receive a food reward, the TSP1/2-KO mice pressed the lever at significantly lower rates than the WT mice (Fig 2C). This suggests that mice lacking TSP1/2 have reduced lever press performance during instrumental tasks requiring more effort, a phenotype that is opposite of α2δ-1 KOs [54].

**Fig 2.**
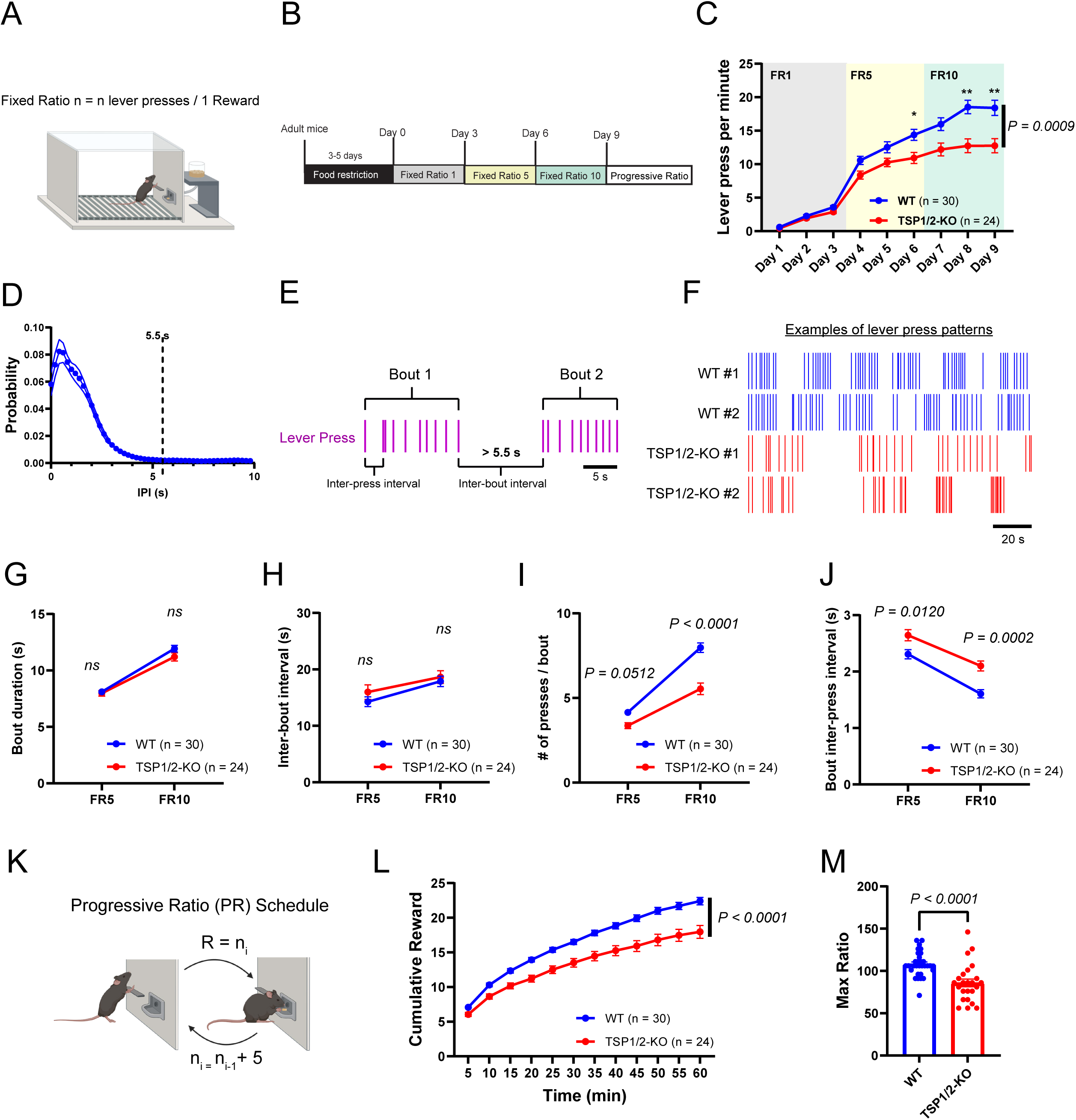
Global loss of TSP1/2 reduces instrumental performance. **(A)** Schematic representation of the Skinner box used for operant training. **(B)** Schematic representation of the training schedule used for the WT and TSP1/2-KO mice. **(C)** Lever press (LP) per minute for the 9 days on the Fixed ratio (FR) schedules for WT (n = 30 mice; 15 males and 15 females) and TSP1/2-KO (n = 24 mice; 12 male and 12 female). Repeated measures Two-way ANOVA. Main effects of Days [F (2.672, 139.0) = 242.9, p < 0.0001], main effect of Genotype [F (1, 52) = 12.31, p = 0.0009], and interaction [F (8, 416) = 7.772, p < 0.0001]. Sidak’s multiple comparisons test with alpha = 0.05. **(D)** Probability of inter-press interval used to identify lever press bouts. **(E)** Schematic representation of lever press per bout, inter-press interval, and inter-bout interval. **(F)** Representative lever press raster plots for FR10 day 9 of LP performance for WT and TSP1/2-KO mice. **(G)** Number of presses per bout. Repeated measures Two-way ANOVA. Main effects of Schedule [F (1, 52) = 380, p < 0.0001], main effect of Genotype [F (1, 52) = 26.84, p < 0.0001], and interaction [F (1, 52) = 28.71, p < 0.0001]. Sidak’s multiple comparisons test with alpha = 0.05. **(H)** Bout duration. Repeated measures Two-way ANOVA. Main effects of Schedule [F (1, 52) = 249.1, p < 0.0001], no effect of Genotype [F (1, 52) = 1.181, p = 0.2823] or interaction [F (1, 52) = 1.245, p = 0.2697]. Sidak’s multiple comparisons test with alpha = 0.05. **(I)** Bout inter-press interval. Repeated measures Two-way ANOVA. Main effects of Schedule [F (1, 52) = 171.8, p < 0.0001], main effect of Genotype [F (1, 52) = 14.23, p = 0.0004], and interaction [F (1, 52) = 2.748, p = 0.1034]. Sidak’s multiple comparisons test with alpha = 0.05. **(J)** Inter-bout interval. Repeated measures Two-way ANOVA. Main effects of Schedule [F (1, 52) = 25.63, p < 0.0001], main effect of Genotype [F (1, 52) = 0.8564, p = 0.3590], and interaction [F (1, 52) = 0.6097, p = 0.4384]. Sidak’s multiple comparisons test with alpha = 0.05. **(K)** Schematic representation of the Progressive Ratio (PR) schedule. **(L)** Cumulative reward obtained during the progressive ratio (PR) task (bin=5 min) for WT (n = 30; 22.40 ± 0.531 rewards); TSP1/2-KO (n = 24; 17.96 ± 0.919 rewards). Repeated measures Two-way ANOVA. Main effects of Genotype [F (1, 52) = 21.48, p < 0.0001] and Time [F (1.485, 77.24) = 677.4, p < 0.0001] and interaction [F (11, 572) = 10.91, p < 0.0001]. **(M)** Quantification of Max Ratio during the PR test. Unpaired Two-tailed T-test. WT (n = 30; 108 ± 2.654) and TSP1/2-KO (n = 24; 85.79 ± 4.5999) [t (52) = 4.383]. Data shown as mean ± s.e.m.

To determine how the reduced performance of TSP1/2-KO mice occurred, we performed a bout analysis during FR5 and FR10 of the instrumental behavior task. To do so, first, we calculated the probability distribution of inter-press intervals (IPI) (Fig 2D) to determine the start and end of each bout (Fig 2E). Then we analyzed the lever press bout patterns of WT and TSP1/2-KO (Fig 2F) mice. The loss of TSP1/2 did not impact bout duration or inter-bout interval (Fig 2G and 2H). However, TSP1/2-KO mice had significantly fewer lever presses per bout during FR10 (Fig 2I) and an increased inter-press interval (IPI) within the bouts during FR5 and FR10 (Fig 2J). In other words, TSP1/2-KO mice had fewer lever presses within a bout because the time between one lever press and the next was significantly longer, thereby leading to a reduction in performance.

To further probe the performance of TSP1/2-KOs in this task, we next utilized a progressive ratio (PR) schedule. During the PR schedule, the number of lever presses necessary to receive one food reward is progressively increased by an increment of 5 (for example, 1 LP for the first reward, 6 LPs for the second, 11 LPs for the third, and so on, as depicted in Fig 2K). This manipulation rapidly increases the effort required to earn a reward, and WT mice usually stop performing the task when the task becomes too demanding. We found that TSP1/2-KO mice earn significantly less rewards during the PR schedule than WT controls (Fig 2L). Thus, TSP1/2-KO mice had a significantly reduced maximum ratio – i.e. the maximum number of lever presses executed for a single reward at the end of the PR task (Fig 2M). These findings reveal that TSP1/2-KO mice have significantly reduced performance during instrumental tasks requiring increased effort. This phenotype is also the opposite of what we previously found in global and circuit-specific α2δ1 KOs [54].

### TSP1/2-KO mice are not hypoactive, have normal food consumption, and can learn action-outcome contingency

The impaired performance of TSP1/2-KO mice during high-effort instrumental tasks could be due to hypoactivity of these mice. To test this possibility, we conducted an open-field test to track and quantify the motor activity of WT and TSP1/2-KO mice in a novel environment (Fig 3A). There were no differences in the total distance traveled (Fig 3B) or the percentage of time spent in the center zone of the open field arena due to loss of TSP1/2 (Fig 3C). These results show that the reduced performance of TSP1/2-KO mice during high-effort instrumental tasks cannot be attributed to overall lower locomotive activity or anxiety-like behaviors.

**Fig 3.**
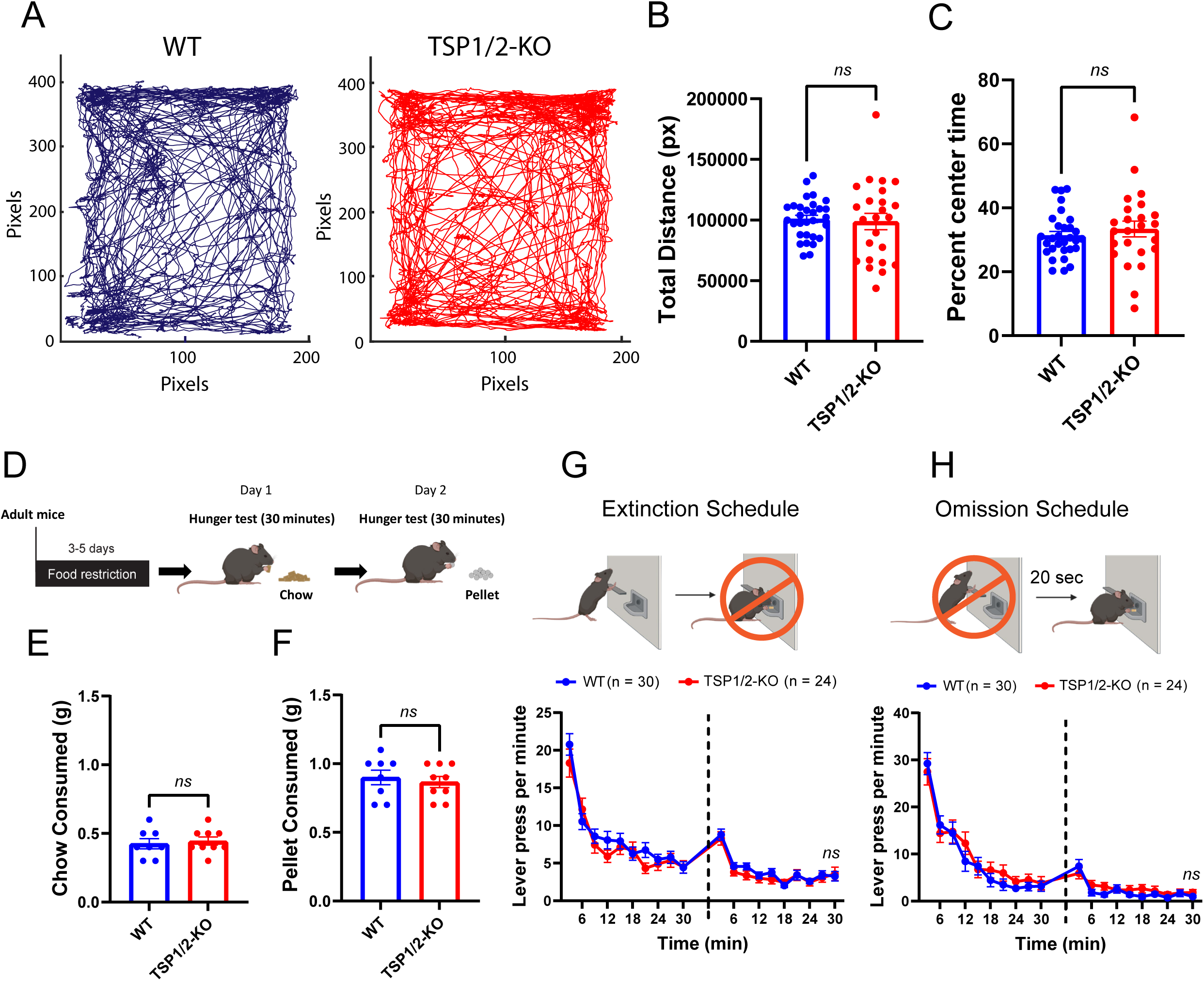
Loss of TSP1/2 does not impact locomotion, hunger levels, or the learning of action-outcome contingency. **(A)** Example of open field patterns of locomotion by WT and TSP1/2-KO mice. **(B)** Bar graph of the total distance traveled in pixels for WT (n = 30; 1.01*10^5^ ± 2.9*10^3^ pixels) and TSP1/2-KO (n = 24; 9.8*10^4^ ± 6.8*10^3^ pixels) mice. Unpaired Two-tailed T-test. [t (31.95) = 0.2708]. **(C)** Percent of time spent in the center of the arena by WT (n = 30; 31.34 ± 1.29%) and TSP1/2-KO (n = 24; 33.37 ± 2.49%) mice. Unpaired Two-tailed T-test. [t (35.04) = 0.7221]. Data shown as mean ± s.e.m. **(D)** Schematic representation of the strategy used to assess hunger levels for WT and TSP1/2-KO mice. **(E)** Quantification of chow consumed by food-restricted WT and TSP1/2-KO mice. Unpaired Two-tailed t-test. WT (0.425 ± 0.037), TSP1/2-KO (0.444 ± 0.029) [t (15) = 0.418]. **(F)** Quantification of food pellets consumed by food-restricted WT and TSP1/2-KO mice. Unpaired Two-tailed t-test. WT (0.90 ± 0.053), TSP1/2-KO (0.867 ± 0.041) [t(15) = 0.502]. WT (n = 8), TSP1/2-KO (n = 9). **(G)** Top: Schematic representation of the extinction schedule. Bottom: Lever press per minute in 3 min bins for the 2 days (dashed line) of testing during the extinction schedule for WT (n =30, 15 male and 15 female) and TSP1/2-KO (n = 24, 12 male and 12 female) animals. Repeated measures Two-way ANOVA. Main effect of Time [F (7.824, 406.9) = 59.40, p < 0.0001] and no effect of genotype [F (1, 52) = 0.7345, p = 0.3954] and interaction [F (19, 988) = 0.9627, p = 0.5039]. **(H)** Top: Schematic representation of the omission schedule. Bottom: Lever press per minute in 3 min bins for the 2 days (dashed line) of testing during the omission schedule. Repeated measures Two-way ANOVA. Main effect of Time [F (6.523, 339.2) = 55.36, p < 0.0001] and no effect of genotype [F (1, 52) = 0.7130, p = 0.4023] and interaction [F (19, 988) = 0.6367, p = 0.8801].

It is possible that TSP1/2-KO mice are less motivated by food, which would mean that they would be satiated with smaller amounts of food compared to WT. To test this, we quantified the amount of home cage food (chow) or reward food (pellets) that food-deprived WT or TSP1/2-KO mice would consume in 30 minutes (Fig 3D). We found that there were no differences in the amount of chow or pellets consumed between WT and TSP1/2-KO mice (Fig 3E and 3F). These results show that when given free access, the TSP1/2-KOs consume equal amounts of food compared to WT, and they consume more pellet (reward food) than home cage chow. Therefore, we conclude that TSP1/2-KOs are not less motivated by food when compared to WT.

Next, we tested whether TSP1/2-KO mice are deficient in learning action-outcome contingency. Such a deficit would explain why, as the contingency changes from FR1 to FR5 or FR5 to FR10, TSP1/2 KOs cannot adapt to changes and execute increased lever presses accordingly. We used extinction and omission schedules to determine if TSP1/2-KO mice can extinguish lever press behavior when the reward is no longer supplied or learn the opposite contingency that lever press delays food reward.

WT mice drastically reduce their lever pressing during the extinction test because it is no longer rewarded. We found that TSP1/2-KO mice extinguished their lever press behavior similarly to WT mice (Fig 3G). During the omission test, reward delivery is delayed every time the mice pressed the lever. The WT mice rapidly reduce their lever press rates. The TSP1/2-KO decreased their lever press rates similarly during the omission test (Fig 3H). These findings show that TSP1/2-KO are not hypoactive, have normal food consumption, and can learn action-outcome contingency like WT mice. Taken together with our earlier findings, we concluded that TSP1/2-KO mice have a specific deficit in performing high-effort tasks. Interestingly, these deficits oppose the phenotypes observed in the global and circuit-specific α2δ1 KOs [54].

### Lack of TSP1/2 does not impair training-induced excitatory synapse formation

We next investigated whether the constitutive loss of TSP1/2 prevents training-induced synapse formation, which was the case in α2δ-1 KO mice [54]. To do so, we quantified intracortical (VGluT1/PSD95 positive) synapses in the ACC of trained WT and TSP1/2-KO mice and compared them to their untrained controls (Fig 4A). Untrained mice were food-restricted as the trained mice but were given free access to the same amount of food rewards in the training chamber without the need to press a lever.

**Fig 4.**
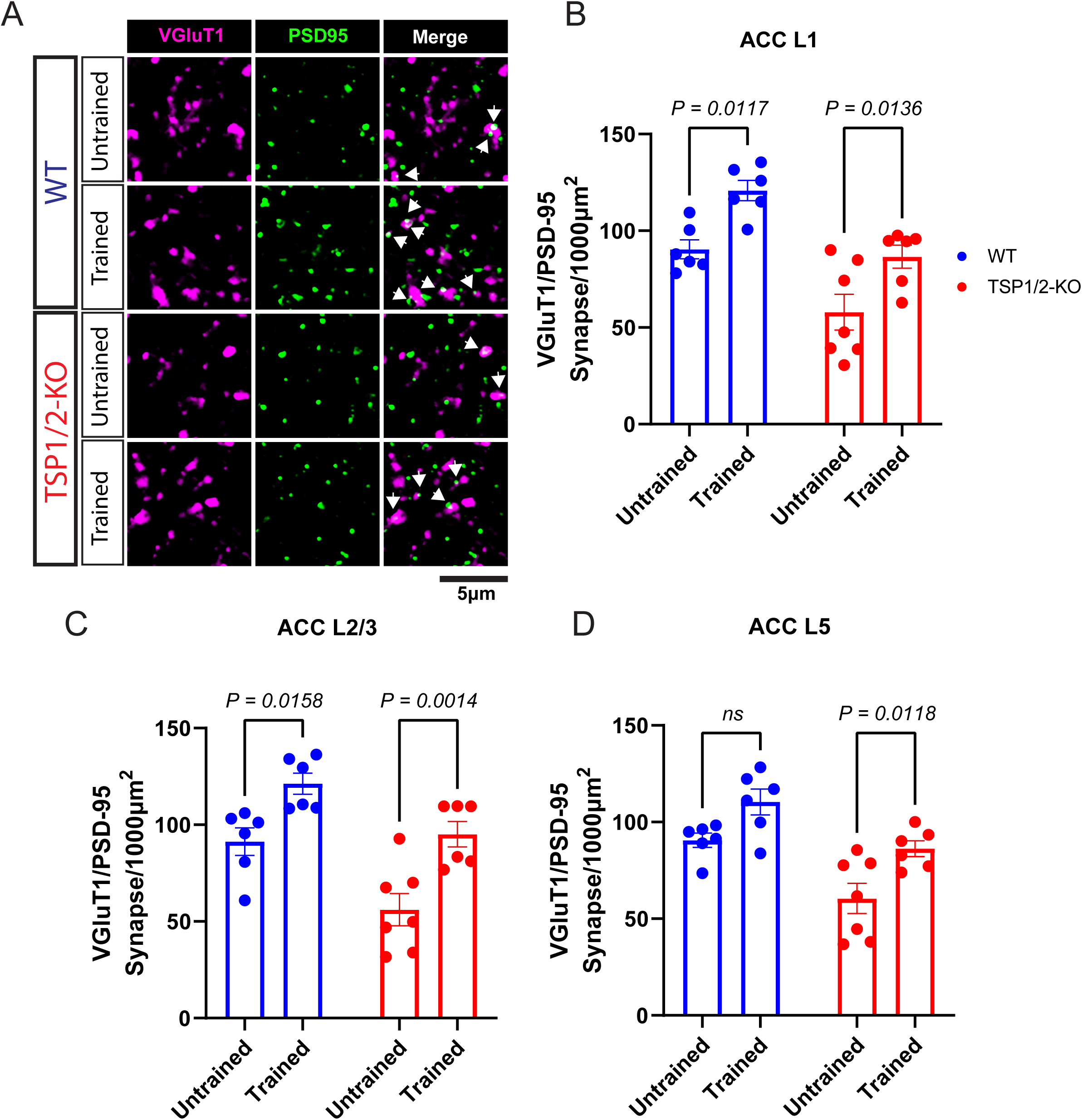
Training-induced excitatory synaptogenesis is not impacted by the loss of TSP1/2. **(A)** Representative images of VGluT1/PSD-95 staining in the ACC of untrained and trained WT and TSP1/2-KO mice. **(B)** Comparisons between ACC L1 untrained and trained WT and TSP1/2-KO mice showing the VGluT1/PSD-95-positive synaptic density. Two-way ANOVA. Main effect of genotype [F (1, 21) = 23.55, p < 0.0001], Training [F (1, 21) = 18.39, p = 0.0003], and no interaction [F (1, 21) = 0.016, p = 0.901]. **(C)** Comparison between ACC L2/3 untrained and trained WT and TSP1/2-KO mice showing the VGluT1/PSD-95-positive synaptic density. Two-way ANOVA. Main effect of genotype [F (1, 21) = 18.72, p = 0.0003], Training [F (1, 21) = 23.60, p < 0.0001], and no interaction [F (1, 21) = 0.401, p = 0.534]. **(D)** Comparison between ACC L5 untrained and trained WT and TSP1/2-KO mice showing the VGluT1/ PSD-95-positive synaptic density. Two-way ANOVA. Main effect of genotype [F (1, 21) = 20.04, p = 0.0002], Training [F (1, 21) = 14.10, p = 0.0012], and no interaction [F (1, 21) = 0.2448, p = 0.626]. (n = 6-7 mice per condition; 3 images per animal). Sidak’s multiple comparisons test with alpha = 0.05 for adjusted p-value. Data shown as mean ± s.e.m.

Training increased excitatory synapse density in WT and TSP1/2-KO ACC layers 1 and 2/3 (Fig 4B and 4C). In ACC L5, like our previous findings [54], training did not significantly increase excitatory synapse density in WT mice (Fig 4D), whereas training also significantly increased excitatory synapse density in the ACC L5 of TSP1/2-KO mice (Fig 4D). These findings indicate that unlike in mice lacking α2δ-1, the loss of TSP1/2 does not fully impair training-induced synapse formation in the ACC. This increase in synapse formation in the ACC likely underlies why TSP1/2-KO mice do not exhibit increased effort exertion in the operant task. These findings also suggest that there are other α2δ-1 ligands, such as other TSP isoforms, which may be upregulated during training and performance of this task.

### Loss of TSP1/2 significantly impacts ACC gene expression

To identify the molecular pathways leading to decreased instrumental performance in the TSP1/2-KO mice, we first investigated the baseline changes in the gene expression of untrained TSP1/2KO mice compared to their WT controls. To do so, we performed RNA sequencing (RNA-seq) from the ACCs of the untrained TSP1/2-KO and WT control mice (Fig 5A). RNA-seq analysis identified 173 differentially expressed genes (DEGs) with a false discovery rate of less than 0.1 (FDR < 0.1) and a fold change of > |1.5| (or log fold change (logFC) > |0.58|). 60 genes were downregulated and 113 were upregulated in the untrained TSP1/2-KO compared to WT (Fig 5B). For downregulated genes analysis of GO terms showed a small but significant enrichment of genes associated with ribonuclease complex and activity, damaged DNA binding, and cell killing (S2A – 2C Fig, S1 File).

**Fig 5.**
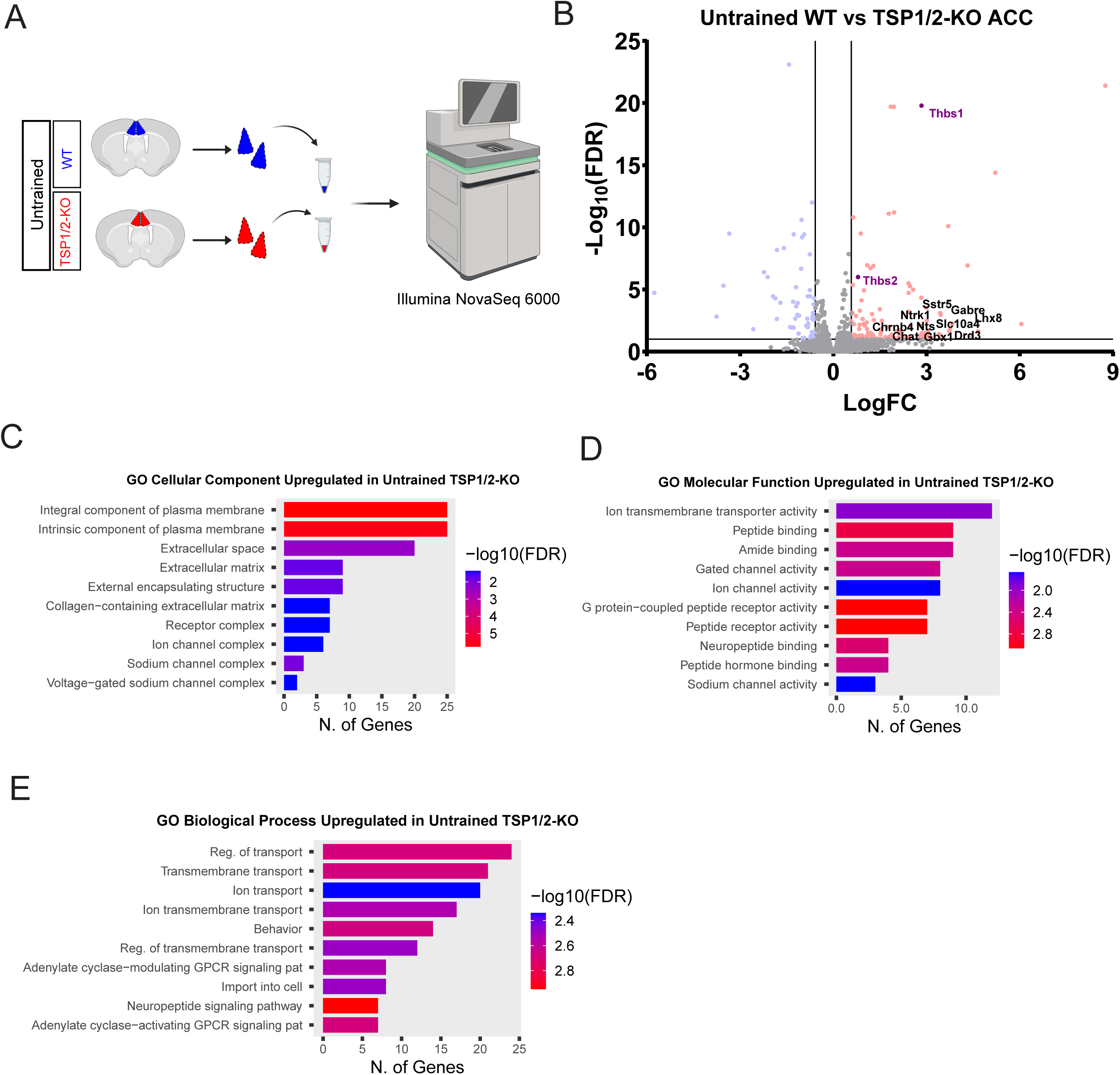
Global loss of TSP1/2 upregulates the expression of genes associated with inhibitory neurons. **(A)** Schematic representation of the ACC microdissection and RNA-seq from untrained WT and untrained TSP1/2-KO mice. **(B)** Volcano plot of differentially expressed genes (DEGs) in the ACC of untrained TSP1/2-KO (n = 8, 4 males, 4 females) compared to WT (n = 8, 4 males, 4 females). Some genes of interest are labeled with the gene symbol. **(C)** Gene Ontology (GO) Cellular Components for genes that are upregulated in untrained TSP1/2-KO compared to untrained WT (ShinyGo 0.80, FDR < 0.05). **(D)** Gene Ontology (GO) Molecular Function for genes that are upregulated in untrained TSP1/2-KO compared to untrained WT (ShinyGo 0.80, FDR < 0.05). **(E)** Gene Ontology (GO) Biological Process for genes that are upregulated in untrained TSP1/2-KO compared to untrained WT (ShinyGo 0.80, FDR < 0.05).

Of the 113 upregulated genes, analysis of gene ontology (GO) terms for cellular components showed enrichment for genes associated with integral membrane proteins and extracellular matrix (Fig 5C, S1 File). This was accompanied by the enrichment of genes associated with molecular functions such as peptide binding and receptor, and ion transmembrane transporter activity (Fig 5D, S1 File), and biological processes such as transmembrane transport and behavior (Fig 5E, S1 File). Taken together, these results showed that in untrained TSP1/2-KO mice, there are significant changes in the expression of genes that are related to neuronal activity and behavior.

Surprisingly, our sequencing data revealed a significant upregulation of *Thbs1* and *Thbs2* mRNAs in the KOs. The TSP1/2-KO mice were generated by excision of exons 2 and 3 in both genes [72,73]. We investigated the abundance of these exons in our sequencing data and found that the Thbs1/2 transcripts we detected did not include reads in exons 2 and 3 (S2D Fig). Furthermore, we performed differential expression at the exon level for *Thbs1* and *Thbs2* and found a significant downregulation of *Thbs1* exon 2 and *Thbs2* exons 2 and 3 (S2 File). These analyses showed that even though Thbs1 and Thbs2 transcripts were significantly upregulated in the TSP1/2 double KOs, the deleted exons in the *Thbs1* or *Thbs2* are appropriately excised and not detected in our RNA sequencing (S2D Fig). These results are in line with previous characterization of these KO mice, showing no TSP1 or 2 proteins are present [63]. However, upregulation of Thbs1 and 2 mRNA, which cannot be translated into intact protein, might trigger compensatory gene expression changes as it was shown for other KOs [74].

Taking a closer look at the DEGs in the untrained TSP1/2-KO compared to untrained WT ACC, we found that among the upregulated genes are a group of genes associated with inhibitory neurons, such as Lhx8, Chat, Ntrk1, Gbx1, and Sstr5 (Fig 5B). LIM Homeobox 8 (Lhx8) is a transcription factor that is critical for the development of interneuron progenitors and cholinergic neurons [75–77], which express choline acetyltransferase (ChAT) [78]. Lhx8 modulates neurotrophic receptor tyrosine kinase 1 (Ntrk1) [79]. In the cortex, a subset of vasoactive intestinal peptide-expressing (VIP+) interneurons can utilize acetylcholine and GABA for synaptic transmission [80,81]. Somatostatin receptor 5 (Sstr5) is another gene upregulated due to the loss of TSP1/2 in the ACC. Sstr5 is highly expressed in parvalbumin (PV), somatostatin (SST), and VIP interneurons, which together account for over 80% of the cortical inhibitory neuron population [82]. Gamma-aminobutyric acid A receptor subunit epsilon (Gabre) is also among the genes upregulated in the ACC of untrained TSP1/2-KO mice (Fig 5B), suggesting an increase in GABAergic transmission. These results show that developmental loss of TSP1/2 alters the expression of genes associated with interneurons and inhibitory synapses.

To investigate how instrumental training changes gene expression in the ACC of TSP1/2-KO mice, we also isolated their ACC after training, extracted RNA, and performed RNA sequencing (Fig 6A). Our analysis identified 344 DEGs with a false discovery rate of less than 0.1 (FDR < 0.1) and a fold change > |1.5| (logFC > |0.58|) (277 downregulated and 67 upregulated genes) between untrained and trained TSP1/2-KO mice (Fig 6A).

**Fig 6.**
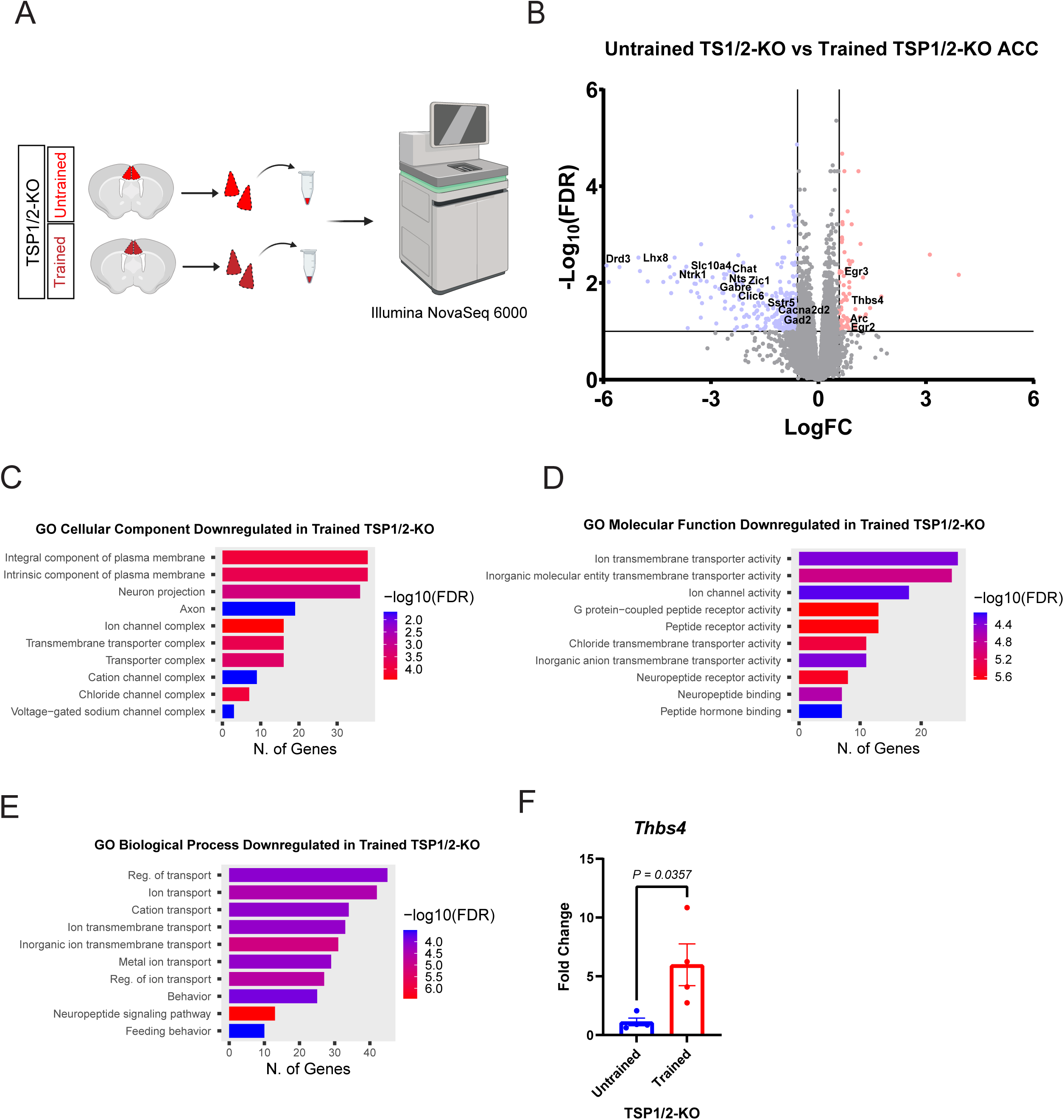
Training downregulates the expression of genes associated with inhibitory neurons and increases *Thbs4* expression in TSP1/2-KO ACC. **(A)** Schematic representation of the ACC microdissection and RNA-seq from untrained TSP1/2-KO and trained TSP1/2-KO mice. **(B)** Volcano plot of DEGs in the ACC (n = 8, 4 males, 4 females) of trained TSP1/2-KO (n = 8, 4 males, 4 females) compared to untrained. Some genes of interest are labeled with the gene symbol. **(C)** Gene Ontology (GO) Cellular Components for genes that are downregulated in trained TSP1/2-KO compared to untrained TSP1/2-KO (ShinyGo 0.80, FDR < 0.05). **(D)** Gene Ontology (GO) Molecular Function for genes that are downregulated in trained TSP1/2-KO compared to untrained TSP1/2-KO (ShinyGo 0.80, FDR < 0.05). **(E)** Gene Ontology (GO) Biological Process for genes that are downregulated in trained TSP1/2-KO compared to untrained TSP1/2-KO (ShinyGo 0.80, FDR < 0.05). **(F)** Quantitative PCR analysis of *Thbs4* fold change in the ACC of TSP1/2-KO mice (n = 4 mice per condition). Unpaired Two-tailed t-test. Untrained (1.114 ± 0.325), Trained (5.977 ± 1.774) [t (6) = 2.697, p = 0.0357]. Data shown as mean ± s.e.m.

GO analysis revealed enrichment of terms for cellular components of the downregulated genes associated with integral membrane proteins, neuron projection, and axon (Fig 6B, S1 File). This was accompanied by enrichment for genes associated with molecular functions related to transmembrane transporter activity, peptide binding, and receptor activity (Fig 6C, S1 File), and biological processes related to ion and transmembrane transport and behavior (Fig 6D, S1 File). Surprisingly, majority of the GO terms for downregulated genes in trained TSP1/2-KO ACC are the terms for upregulated genes in untrained TSP1/2-KO compared to untrained WT (Fig 5C-5E). These results suggest that training reverses some of the gene expression profiles of TSP1/2-KO ACC.

On the other hand, analysis of GO terms for upregulated genes shows enrichment of genes associated with the cell surface, endoplasmic reticulum lumen, signaling receptor activity, protein folding, and cellular stress response (S2E-2G Fig). The enrichment of these GO terms might be due to the expression of *Thbs1* and *Thbs2* mRNAs in the absence of exons 2 and 3 (S2D Fig). This may have impaired or dysregulated the production of their corresponding proteins, leading to the upregulation of genes associated with protein folding.

Interestingly, training also upregulated the expression of thrombospondin isoform 4 (TSP4), which promotes synapse formation and neurite outgrowth [43,49]. We confirmed this RNA-seq result by qPCR analysis and found that *Thbs4* (TSP4) is indeed upregulated in the trained compared to the untrained ACC of TSP1/2-KO mice (Fig 6E). This result shows that in the absence of TSP1/2, TSP4 expression is upregulated after training and suggests that upregulated TSP4 expression may trigger the training-induced synaptogenesis in the ACC or the TSP1/2-KO mice.

In agreement with this, similar to our previous finding for trained WT ACC [54], training in TSP1/2-KO mice upregulated immediate early genes such as Arc, Egr2, and Egr3 (Fig 6A), which play a role in synaptic plasticity and memory-related processes [83–85]. Interestingly, the training downregulated the expression of genes such as Lhx8, Chat, Ntrk1, Gabre, Sstr5, and glutamic acid decarboxylase 2 (Gad2), which are associated with interneurons (Fig 6A). Gad2 encodes the protein glutamate decarboxylase 65 (GAD65) which synthesizes glutamate and is a common marker for inhibitory neurons [86]. These results strongly suggest that gene expression related to inhibitory neurons and synapses are down-regulated in the trained TSP1/2-KO mice compared to untrained KOs. This is the opposite of what we have observed when we compared the untrained WT and TSP1/2-KO mice (Fig 5B).

To implicate which cell types may be impacted in the trained versus untrained TSP1/2-KO ACC, we compared our DEGs to previously published cortical single cell dataset [87]. To do so we used the single-cell mapper package in R [88] to generate cell-type specificity scores using the downregulated DEGs in trained TSP1/2-KO ACC compared to untrained. We found the DEGs downregulated in the trained versus untrained TSP1/2-KO ACCs to be significantly enriched for genes associated with interneurons and GABAergic neurons (p = 0.036) (S3 File). Taken together, the results of our RNA-seq analyses suggest that genes linked to inhibitory neurons and synapses are upregulated in the ACC of untrained TSP1/2-KO mice compared to WT, and training downregulates the genes associated with inhibitory neurons in the ACC of TSP1/2-KO mice. On the other hand, training upregulates expression of Thbs4, a synaptogenic isoform of TSP1 and 2. This upregulation is likely causing the training-induced synapse formation in these KO mice.

### TSP1/2 KO mice have increased inhibitory synapse numbers and function in the ACC

The role of TSPs in inhibitory synapse formation *in vivo* is unknown. Given our findings that genes associated with interneurons are upregulated in the ACC of TSP1/2-KO mice compared to WT (Fig 5), we investigated whether the loss of TSP1/2 alters inhibitory synapse density in the ACC. To do this, we quantified the colocalization of the inhibitory presynaptic marker, VGAT, and post-synaptic marker, Gephyrin, in the ACC of adult WT and TSP1/2-KO mice (Fig 7A). Interestingly, we found a strong and significant increase in inhibitory synapse density in the ACC layers 1, 2/3, and 5 of TSP1/2-KO mice compared to WT (Fig 7B-7D).

**Fig 7.**
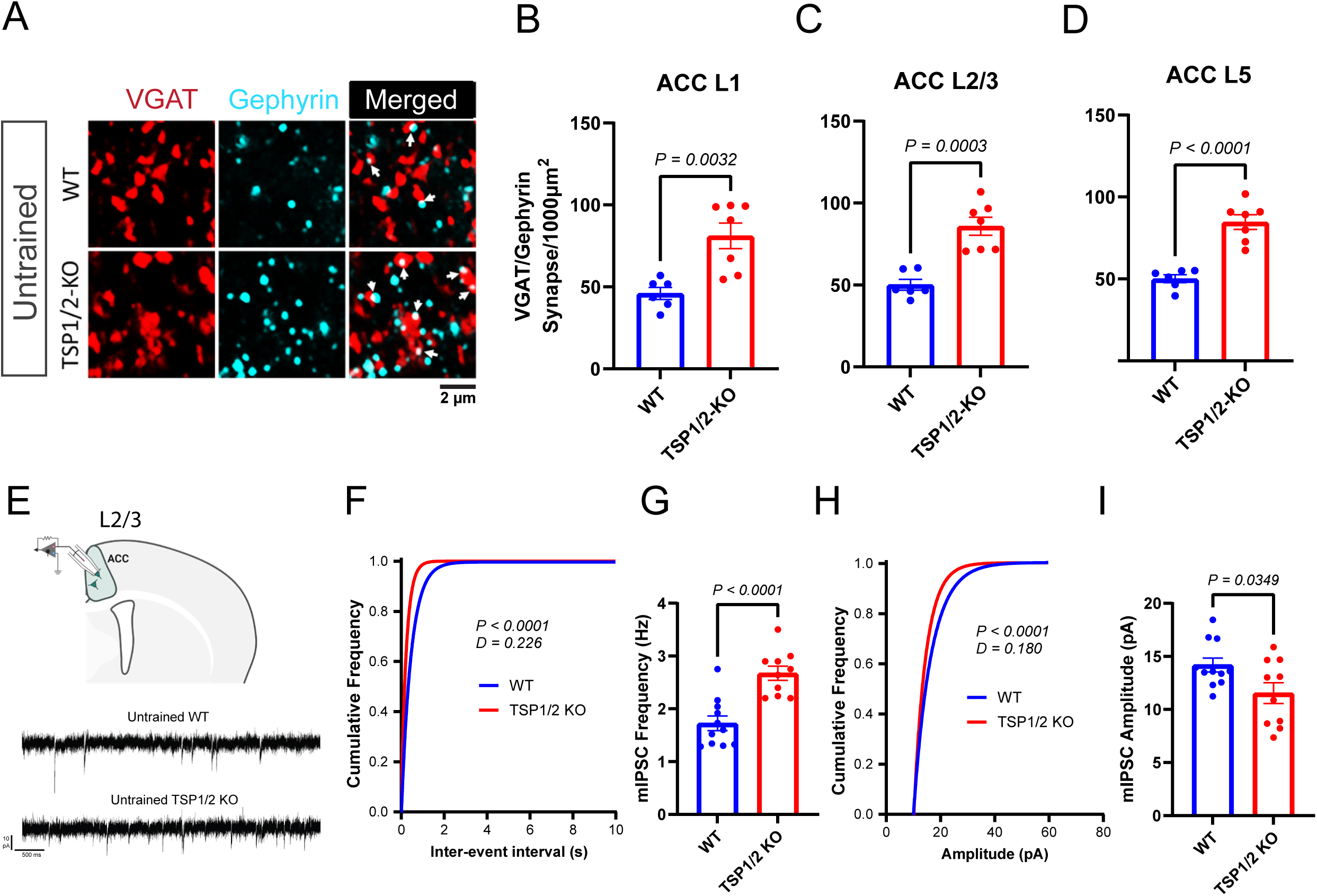
Inhibitory synapse density is enhanced in the ACC of TSP1/2-KOs. **(A)** Top: Representative images of VGAT/Gephyrin staining in the anterior cingulate cortex of untrained WT and TSP1/2-KO mice. **(B)** Quantification of VGAT/Gephyrin co-localized puncta in ACC L1 of untrained WT (46.05 ± 3.68), TSP1/2-KO (81.08 ± 7.80). Unpaired Two-tailed t-test. [t (8.468) = 4.061, p = 0.0032]. **(C)** VGlut1/PSD-95 co-localized puncta in ACC L2/3. WT (50.17 ± 3.287), TSP1/2-KO (85.83 ± 5.493) Unpaired Two-tailed t-test. [t (9.606) = 5.566, p = 0.0003]. **(D)** VGlut1/PSD-95 co-localized puncta in ACC L5. WT (50.10 ± 2.454), TSP1/2-KO (84.69 ± 4.519) Unpaired Two-tailed t-test. [t (9.111) = 6.726, p < 0.0001]. **(E)** Top: Schematic representation of the electrophysiological recordings from ACC L2/3 neurons. Bottom: Example traces from mIPSC recordings from untrained WT and TSP1/2-KO mice. **(F)** Cumulative distribution of the inter-event interval of mIPSC in WT (n = 2 mice; 11 cells) and TSP1/2-KO (n = 3 mice; 10 cells) mice. Kolmogorov-Smirnov test [D = 0.226, p < 0.0001]. **(G)** Average frequency of mIPSC in WT (n = 2 mice, 11 cells; 1.725 ± 0.139 Hz) and TSP1/2-KO (n = 3 mice, 10 cells; 2.678 ± 0.134 Hz) mice. Unpaired t-test [t (19) = 4.899, p < 0.0001]. **(H)** Cumulative distribution of the amplitude of mIPSC in pA for both WT (n = 2 mice; 11 cells) and TSP1/2-KO (n = 3 mice; 10 cells) mice. Kolmogorov-Smirnov test [D = 0.180, p < 0.0001]. **(I)** Average amplitude of mIPSC in WT (n = 2 mice, 11 cells; 14.19 ± 0.659 pA) and TSP1/2-KO (n = 3 mice, 10 cells; 11.54 ± 0.984 pA) mice. Unpaired t-test [t (19) = 2.272, p = 0.0349].

To determine whether this increase in inhibitory synapse structures reflects an increase in inhibitory synapse function, we performed mIPSC recordings on neurons in layers 2/3 of the ACC (Fig 7E). Indeed, we found a significant increase in the frequency of mIPSCs (Fig 7F and 7G) which aligns with the observed increase in synapse numbers. Interestingly, this increase in frequency was accompanied by a significant decrease in the amplitude (Fig 7H and 7I), of mIPSCs in L2/3 of TSP1/2-KO ACC compared to WT. The decrease in mIPSC amplitude could be due to compensatory mechanisms or homeostatic plasticity causing changes to the amount of GABA released from a single vesicle [89]. Alternatively, ACC L2/3 neurons in the TSP1/2-KO mice may have a reduced number of post-synaptic GABAA receptors clustered at synapses [90]. The increase in inhibitory synapses (Fig 7A-7I) and the decrease in excitatory synapses (Fig 1) suggest a disruption in excitation/inhibition balance in the ACC of the untrained TSP1/2-KO mice.

### Training decreases inhibitory synapse density in the ACC of TSP1/2-KO mice without affecting GABAergic neuron numbers

Operant training does not alter inhibitory synapses in the ACC of WT mice [54]. However, given the surprising result that inhibitory neuron-associated genes are high in untrained TSP1/2-KOs and training diminishes their expression in these mice (Fig 6A), we next tested how inhibitory synapse density is impacted in the ACC of trained TSP1/2-KO mice. To do so, we quantified VGAT/Gephyrin-positive inhibitory synapses in the ACC of trained WT and TSP1/2-KO mice and compared these to their respective untrained controls (Fig 8A-8D). As expected from previous results [54], we found no difference in inhibitory synapse density in layers 1, 2/3, and 5 of the ACC between untrained and trained WT mice (Fig 8B-8D). However, there was a significant decrease in the numbers of inhibitory synapses in the ACC of trained TSP1/2-KOs compared to their untrained controls (Fig 8B-8D). This result revealed that training significantly decreases the number of inhibitory synapses in the ACC of TSP1/2-KO mice. This result aligns with our findings in the RNA-seq experiments on untrained versus trained TSP1/2-KO ACCs, which revealed a reduction in the genes associated with inhibitory neurons and synapses (Fig 6).

**Fig 8.**
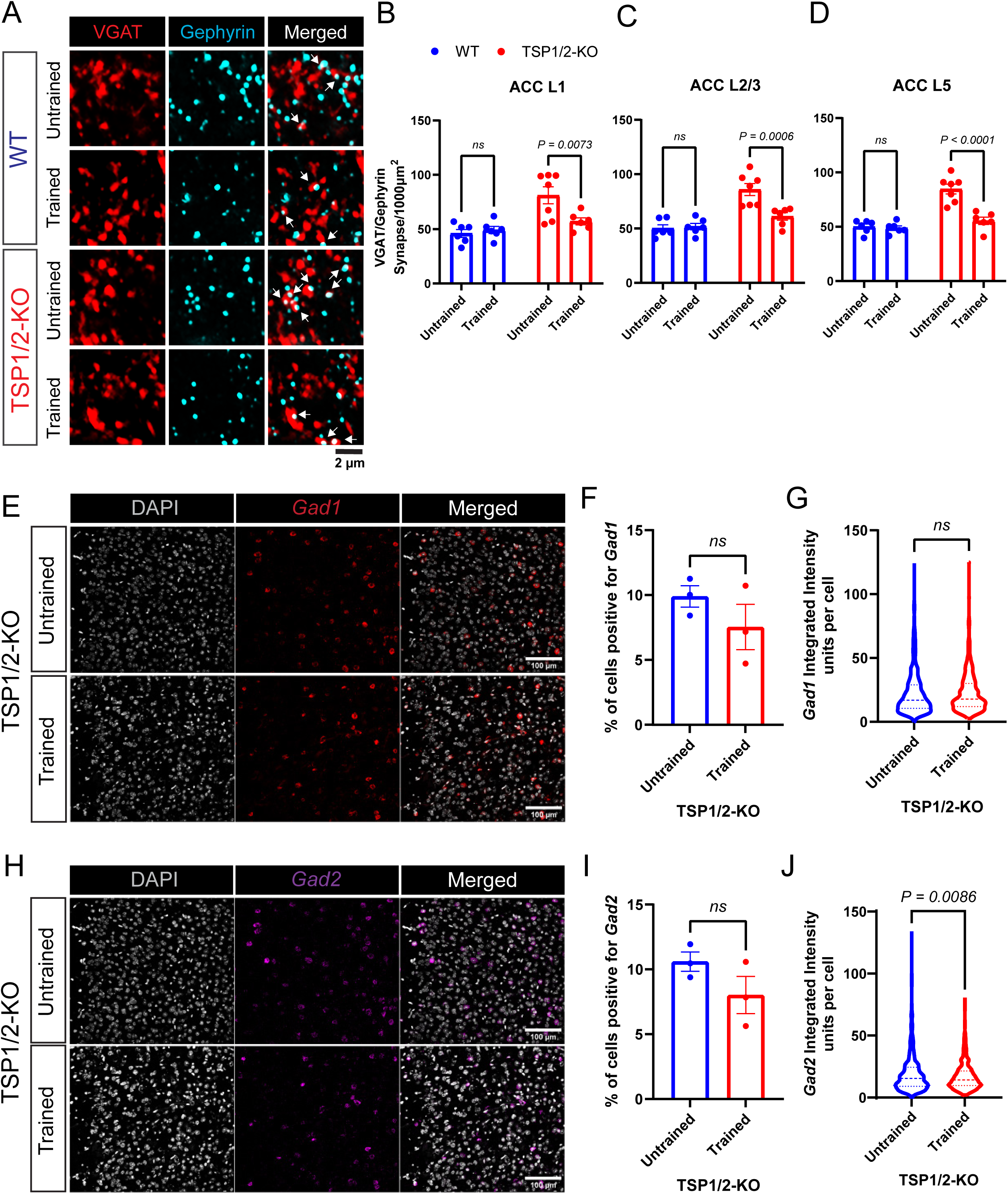
Training reduces inhibitory synapse density in the ACC of TSP1/2-KOs. **(A)** Representative images of VGAT/Gephyrin staining in the anterior cingulate cortex of untrained and trained WT, and TSP1/2-KO mice. **(B)** Comparison between ACC L1 untrained and trained WT and TSP1/2-KO mice showing the VGAT/Gephyrin-positive synaptic density. Two-way ANOVA. Main effect of genotype [F (1, 21) = 16.77, p = 0.0005], no effect of Training [F (1, 21) = 3.87, p = 0.063], and interaction [F (1, 21) = 6.615, p = 0.018]. **(C)** Comparison between ACC L2/3 untrained and trained WT and TSP1/2-KO mice showing the VGAT/Gephyrin-positive synaptic density. Two-way ANOVA. Main effect of genotype [F (1, 21) = 29.84, p < 0.0001], Training [F (1, 21) = 7.91, p = 0.01], and interaction [F (1, 21) = 10.21, p = 0.004]. Sidak’s multiple comparisons test with alpha = 0.05 for adjusted p-value. **(D)** Comparison between ACC L5 untrained and trained WT and TSP1/2-KO mice showing the VGAT/Gephyrin-positive synaptic density. Two-way ANOVA. Main effect of genotype [F (1, 21) = 36.94, p < 0.0001], Training [F (1, 21) = 20.19, p = 0.0002], and no interaction [F (1, 21) = 16.89, p = 0.0005]. (n = 6-7 mice per condition; 3 images per animal). Sidak’s multiple comparisons test with alpha = 0.05 for adjusted p-value. **(E)** Representative images of *Gad1* RNA-FISH in untrained and trained TSP1/2-KO ACC. Scale bar - 100µm. **(F)** Quantification of the percentage of cells (DAPI) positive for *Gad1* in the ACC of TSP1/2-KO mice (n = 3 mice per condition; 3-4 images per mouse). Unpaired Two-tailed t-test. Untrained (9.89 ± 0.821), Trained (7.54 ± 1.741) [t (4) = 1.225, p = 0.2877]. **(G)** Quantification of *Gad1* integrated intensity per cell in the ACC of TSP1/2-KO mice (n = 3 mice per condition; Untrained – 651 cells, Trained – 491 cells). Unpaired Two-tailed t-test. Untrained (22.25 ± 0.661), Trained (22.66 ± 0.684) [t (1140) = 0.424, p = 0.672]. **(H)** Representative images of *Gad2* RNA-FISH in untrained and trained TSP1/2-KO ACC. Scale bar - 100µm. **(I)** Quantification of the percentage of cells (DAPI) positive for *Gad2* in the ACC of TSP1/2-KO mice (n = 3 mice per condition; 3-4 images per mouse). Unpaired Two-tailed t-test. Untrained (10.60 ± 0.740), Trained (8.02 ± 1.429) [t (4) = 1.603, p = 0.1841]. **(J)** Quantification of *Gad2* integrated intensity per cell in the ACC of TSP1/2-KO mice (n = 3 mice per condition; Untrained – 698 cells, Trained – 722 cells). Unpaired Two-tailed t-test. Untrained (19.40 ± 0.603), Trained (17.23 ± 0.506) [t (1218) = 2.631]. Data shown as mean ± s.e.m.

To determine if training decreased the numbers of inhibitory synapses by changing overall inhibitory neuron numbers in the ACC, we performed RNA fluorescent in situ hybridization (RNA-FISH) with probes for *Gad1* (Fig 8E-8G) and *Gad2* (Fig 8H-8J), which are markers of inhibitory neurons [91,92]. We found no difference in the percentage of cells labeled with *Gad1* (Fig 8F) or *Gad2* (Fig 8I) between trained and untrained TSP1/2-KO ACC. This result shows that the inhibitory neuron numbers are not decreased by training in the TSP1/2KO mice.

Quantification of integrated signal intensity per cell showed no difference in *Gad1* (Fig 8G) but a reduction in *Gad2* (Fig 8J) in the trained compared to untrained TSP1/2-KO ACC. The decline in the intensity of Gad2 signal is in line with our RNA-seq findings showing a significant decrease in the expression of *Gad2* mRNA. These data show that instrumental training does not decrease inhibitory cell numbers but downregulates inhibitory neuron connectivity in the ACC of TSP1/2-KO mice. Our findings indicate that training increases excitation and decreases inhibition in the TSP1/2-KO ACC.

### Conditional deletion of TSP1/2 in adult astrocytes does not impair instrumental learning or performance

Thbs1 and Thbs2 are expressed in tissues throughout the body [47,93]. In the brain, astrocytes express Thbs1 and Thbs2 primarily during early postnatal development [43,94,95].

Because constitutive loss of TSP1/2 impaired performance during high-effort instrumental tasks, we tested whether this behavior deficit can be attributed to the loss of TSP1/2, specifically in adult astrocytes. To ablate TSP1 and TSP2 in astrocytes, we utilized a viral approach involving adeno-associated viruses (AAVs) [96]. This astrocyte-specific CRISPR-based gene-editing tool called GEARBOCS (Gene Editing in AstRocytes Based On CRISPR/HITI System) uses a single AAV vector to knock out a gene in mice expressing Cas9 in a Cre recombinase-dependent manner [96]. Using this methodology, we generated AAV vectors carrying guide RNAs (sgRNA) targeting *Thbs1* or *Thbs2* (S3A and S3B Fig) and Cre expressed under the control of the GFABC-1D promoter. As a control, we used the same vector without sgRNAs (S3B Fig).

To eliminate TSP1 and TSP2 expression in adult astrocytes, we injected these AAVs retro-orbitally into 8 to10 week-old adult Cas9-EGFP (loxP-STOP-loxP) [97] mice to specifically knockout both genes in astrocytes (Fig 9A). After three weeks, brain tissues from injected mice were stained with anti-GFP antibodies to determine whether there was Cas9 expression in astrocytes. We found GFP-positive astrocytes throughout the brain with robust labeling in the ACC (Fig 9B). To determine whether this strategy had disrupted *Thbs1* and *Thbs2* genes at the location targeted by the sgRNA, we collected brains from mice injected with the knockout and control AAV vectors. We dissociated the brains, sorted GFP-positive cells, and extracted DNA from these cells. Subsequently, we amplified the targeted regions with PCR and sequenced the amplicons using Sanger sequencing (S3C Fig). The sequencing data revealed indels at the targeted *Thbs1* and *Thbs2* gene loci of extracted DNA from TSP1 and TSP2 conditional knockout (TSP1/2-cKO) GFP-positive astrocytes (S3D and S3E Fig). These indels cause a frameshift and premature STOP codons in the coding sequence.

**Fig 9.**
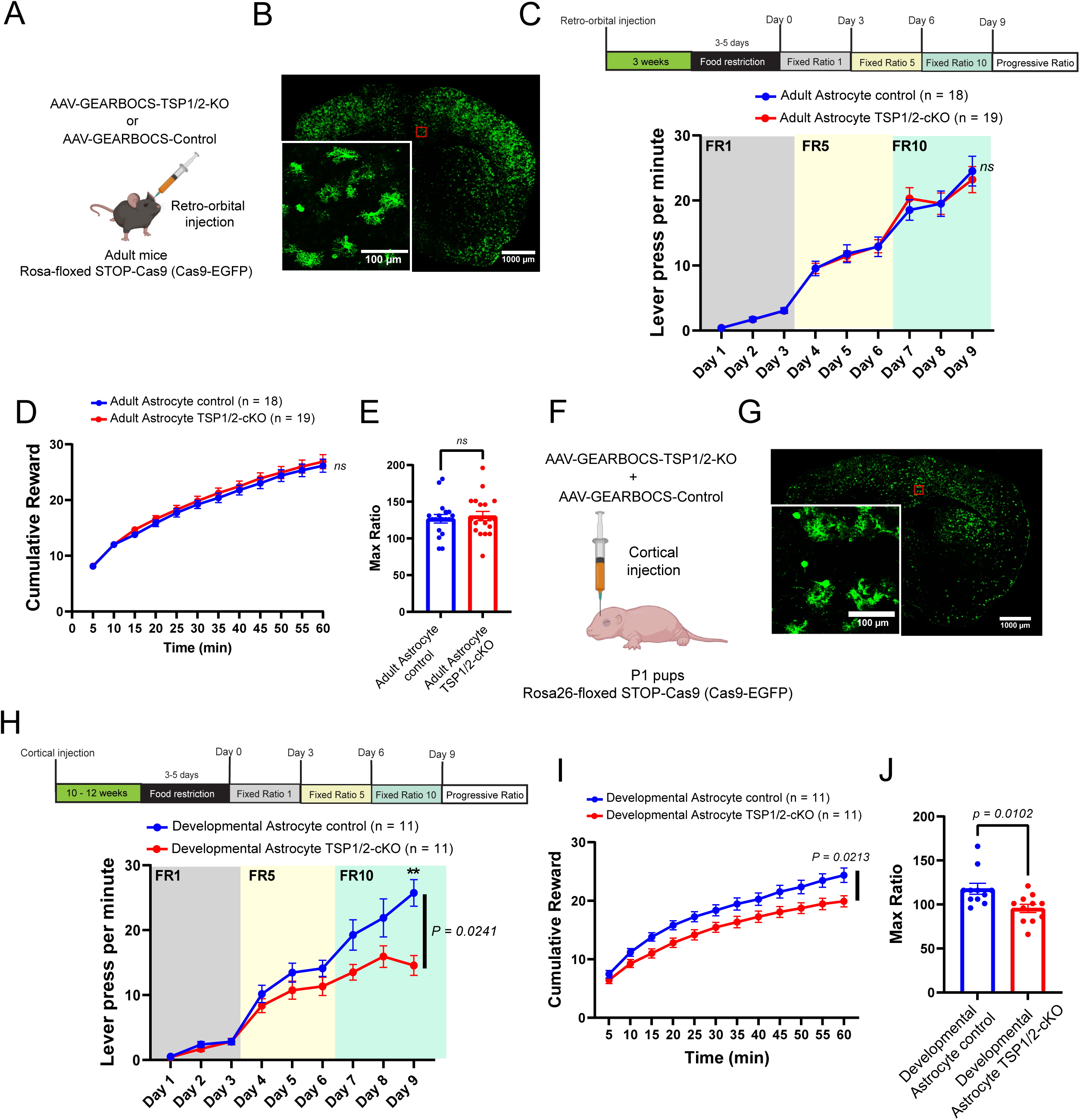
Loss of TSP1/2 in developing but not adult astrocytes reduces instrumental performance. **(A)** Schematic representation of the retro-orbital injection of AAV vector to ablate TSP1/2 in adult astrocyte using a CRISPR-based approach. **(B)** Tile scan image of a coronal brain section from a Cas9-EGFP mouse retro-orbitally injected with AAV vector to ablate TSP1/2 in adult astrocyte. Inset - The GPF labeling shows the expression of Cas9 in astrocytes. **(C)** Top: Schematic representation of the timeline treatments and training schedules used for the adult control and TSP1/2-cKO mice. Bottom: Lever press (LP) per minute for the 9 days on the FR schedule for control (n = 18, 11 males and 7 females) and TSP1/2-cKO (n = 19, 11 males and 8 females) mice. Repeated measures (RM) Two-way ANOVA. Main effects of Days [F (8, 280) = 155.8, p < 0.0001], no effect of Genotype [F (1, 35) = 0.00023, p = 0.988] nor interaction [F (8, 280) = 0.339, p = 0.95]. Sidak’s multiple comparisons test with alpha = 0.05 for adjusted p-value. **(D)** Cumulative reward count over the PR session (bin=5min) for adult control (n = 18 mice) and TSP1/2-cKO (n = 19 mice). RM Two-way ANOVA. Main effects of Time [F (1.330, 46.54) = 408.5, p < 0.0001], no effect of Genotype [F (1, 35) = 0.308, p = 0.5827] nor interaction [F (11, 385) = 0.274, p = 0.990]. **(E)** Max Ratio for control (n = 18; Max Ratio = 126.8 ± 5.85) and TSP1/2-cKO (n = 19; Max Ratio = 130.5 ± 6.24) animals. Unpaired two-tailed t-test [t (35) = 0.425, p=0.6736]. **(F)** Schematic representation of the cortical injection of AAV vector to ablate TSP1/2 in developing neonatal astrocytes using a CRISPR-based approach. **(G)** Tile scan image of a coronal brain section from a Cas9-EGFP mouse pup intracortically injected with AAV vector to ablate TSP1/2 in developing astrocyte. Inset - The GPF labeling shows the expression of Cas9 in astrocytes. **(H)** Top: Schematic representation of the timeline treatments and training schedules used for the adult control and TSP1/2-cKO mice. Bottom: Lever press (LP) per minute for the 9 days on the FR schedule for developmental control (n = 11, 5 male and 6 female) and TSP1/2-cKO (n = 11, 5 male and 6 female) mice. Repeated measures Two-way ANOVA. Main effect of Days [F (3.098, 61.95) = 96.28, p < 0.0001], and Genotype [F (1, 20) = 5.953, p = 0.0241], and effect of interaction [F (8, 160) = 5.819, p < 0.0001]. Sidak’s multiple comparisons test with alpha = 0.05 for adjusted p-value. **(I)** Cumulative reward count over the PR session (bin=5min) for Developmental control (n = 11) and TSP1/2-cKO (n = 11) mice. Repeated measures Two-way ANOVA. Main effects of Time [F (1.939, 38.79) = 405.6, p < 0.0001], and Genotype [F (1, 20) = 6.241, p = 0.0213], and interaction [F (11, 220) = 3.733, p < 0.0001]. **(J)** Max Ratio for Developmental control (n = 11; Max Ratio = 117.8 ± 6.189) and TSP1/2-cKO (n = 11; Max Ratio = 95.55 ± 4.741) animals. Unpaired two-tailed t-test [t (18.73) = 2.857, p=0.0102]. Data shown as mean ± s.e.m.

To determine whether the CRISPR-mediated loss of TSP1/2 in adult astrocytes impairs goal-directed actions during high effort, we trained adult TSP1/2-cKO and control mice on instrumental tasks as previously described. Quantification of lever pressing showed that there are no differences between control and TSP1/2-cKO mice (Fig 9C).

The specific ablation of TSP1/2 in the adult astrocytes may necessitate a much higher effort requirement during an instrumental task to observe a difference between the control and TSP1/2-cKO mice. To address this, we tested the mice on a progressive ratio (PR) schedule, which increases the number of lever presses for subsequent reward by 5 (Fig 2K). Still, there were no differences in the cumulative reward obtained between the GEARBOCS TSP1/2KO or control-injected mice at the end of the PR session (Fig 9D). There was also no difference in the maximum number of lever presses executed for a food reward (max ratio) between the conditions (Fig 9E). These results show that loss of TSP1/2 in adult astrocytes is not sufficient to impair GDA performance during high effort tasks.

To determine whether the conditional ablation of TSP1/2 in astrocytes impairs goal-directed actions, such as adaptability to changes in action-outcome contingency, we also tested control and TSP1/2-cKO mice on extinction and omission schedules. There were no differences between the control and TSP1/2-cKO mice’s ability to extinguish their lever presses when a reward was no longer delivered (S4A and S4B Fig). Further, the ability to reduce lever presses to avoid a delay in food reward delivery was not impaired by the loss of TSP1/2 in adult astrocytes (S4C and S4D Fig). We also observed no differences in the total distance traveled and percentage of time spent in the center zone of an open-field arena (S4E-4G Fig), indicating no significant impairments to locomotive activity. All these results indicate that astrocytic TSP1/2 expression in adults is not necessary for the learning and performance of instrumental tasks.

### Conditional ablation of TSP1/2 in developing astrocytes impairs performance during high-effort tasks

Given that TSP1/2 ablation in adult astrocytes did not produce any impairment in instrumental operant task performance (Fig 9A-9E), we considered the possibility that astrocytic TSP1/2 signaling is specifically needed during early development for the establishment of the synaptic connections involved in the control of goal-directed actions. In agreement, expression of TSP1 and TSP2 in the cortex is highest during the first postnatal week of development compared to adults [29,94,98,99]. Postnatal protein expression of cortical TSP1/2 is highest during the first week and mostly absent by postnatal day 21 (P21) [29]. Therefore, we hypothesized that the role of astrocytic TSP1/2 is primarily critical during development, and its ablation in neonates will impair goal-directed action performance in adulthood. To test this hypothesis, we injected the AAV vectors to ablate astrocytic TSP1/2 into the neonatal cortex at postnatal day 1 (P1) (Fig 9F). Upon reaching adulthood, brain tissues from injected mice were stained with anti-GFP antibodies to determine whether there was Cas9 expression in astrocytes. We found GFP-positive astrocytes throughout the brain with robust labeling in the cortex (Fig 9G).

The adult mice with developmental TSP1/2-cKO and control astrocytes were trained on instrumental tasks using previously described fixed ratio schedules (Fig 9H). We found that developmental knockout of TSP1/2 in astrocytes impaired instrumental task performance in adults during FR10 (Fig 9H), which is the same phenotype as the global TSP1/2-KOs. There was a significant reduction in lever press rate by the TSP1/2-cKO mice on the last day of FR10 compared to the control mice (Fig 9H). When the mice were tested on the PR task, we also found a significant reduction in cumulative reward in adult mice with developmental astrocyte TSP1/2-cKO compared to control (Fig 9I). Analysis of the maximum number of lever presses executed for a food reward (max ratio) during the PR session shows a significant decrease in the max ratio by developmental astrocyte TSP1/2-cKO mice compared to its virus-injected controls (Fig. 9J).

We also tested control and TSP1/2-cKO mice on extinction and omission schedules to determine if the ablation of TSP1/2 in neonatal astrocytes impairs the learning and adaptability to action-outcome contingency in adulthood. We found no differences between adult mice with developmental TSP1/2-cKO and control astrocytes during the extinction or omission schedule behavior paradigm (S4H-S4K Fig). Interestingly, the open-field test (S4L Fig) showed a reduction in the total distance traveled in the open-field arena by adult mice with developmental astrocyte TSP1/2-cKO compared to the control (S4M Fig). However, there were no differences in the percentage of time spent in the center zone of the arena by the developmental astrocyte TSP1/2-cKO and control mice (S4N Fig). Overall, the data suggests that the loss of TSP1/2 in developing astrocytes impaired instrumental performance during high-effort tasks, a phenotype similar to that of the TSP1/2 global KO mice. Taken together these experiments reveal that expression of TSP1/2 by astrocytes during development is critical for proper performance of goal directed behaviors in adulthood.

## Discussion

Astrocytes promote excitatory synapse formation through the secretion of synaptogenic proteins [26–29]. Thrombospondins are a family of glycoproteins secreted by astrocytes to induce excitatory synaptogenesis through their interaction with the neuronal receptor α2δ-1 [43,51]. The loss of α2δ-1 impairs operant training-induced excitatory synapse formation in the ACC and increases effort exertion during demanding tasks [54]. Here, we show that the loss of two astrocyte-secreted α2δ-1 ligands, thrombospondins 1 and 2 (TSP1/2-KO), do not impair training-induced excitatory synaptogenesis and significantly decreases performance during high-effort tasks, a phenotype opposite of α2δ-1 KOs. Additionally, we found a strong increase in inhibitory synapse number and function in the TSP1/2-KOs and these synapses were eliminated due to operant training. Finally, using cell-type-specific CRISPR/Cas9 approaches, we found that astrocyte-specific ablation of TSP1/2 in early postnatal but not adult astrocytes recapitulates the impaired performance observed in the global TSP1/2-KO mice. Our findings show that astrocyte-secreted synaptogenic TSP1/2 are required during early development to establish correct excitatory and inhibitory connectivity in the ACC for proper control of goal-directed action performance in adulthood.

Studies examining thrombospondins’ role in complex animal behaviors are rare. A recent study showed that activation of Gi protein-coupled receptors in striatal astrocytes increases *Thbs1* mRNA and causes hyperactivity with disrupted attention [53]. Furthermore, blocking the TSP-receptor α2δ-1 function with its specific ligand, gabapentin, was able to reverse the behavior phenotype [53]. However, it is unknown whether the loss of TSP1/2 protein impacts goal-directed behaviors and effort control.

Here, by using the lever-press-for food instrumental training paradigm and global and conditional TSP1/2-KOs, we show a role for astrocyte-derived TSP1/2 in effort control. TSP1/2-KO mice perform less lever presses than the WT mice by decreasing the number of lever presses per bout and increasing their inter-press interval. These changes occur without an increase in bout duration or inter-bout intervals, indicative of a reduction in effort compared to the WT. Previously, lesion of the ACC was shown to shift preference towards low-effort tasks [100,101]. However, we found that increased activity of a particular projection from the ACC to the dorsal medial striatum (DMS) diminishes effort, whereas inhibiting the activity of the same circuit enhances effort exertion [54]. Therefore, an enhancement in the excitation of the ACC to DMS connections may underlie the behavioral phenotypes we observed in TSP1/2-KOs. However, disconnecting the ACC and the basolateral amygdala (BLA) or the ACC and the nucleus accumbens (NAc) shifts rodents towards rewards requiring low effort [102,103], suggesting that ACC projections to these regions within the basal ganglia circuit may also be affected in the TSP1/2-KOs which underlie the lower effort phenotype we observed during instrumental tasks.

Given the importance of TSP-α2δ-1 signaling in excitatory synaptogenesis, we predicted that the constitutive loss of TSP1/2 would phenocopy the maladaptive effort exertion we discovered in the α2δ-1 deficient mice. Instead, we found that unlike mice lacking α2δ-1, knockout of TSP1/2 did not increase effort exertion but decreased performance during high-effort tasks as early as FR5. These opposing phenotypes in the receptor (α2δ-1) versus ligand (TSP1/2) KOs raised several alternative hypotheses for how TSP/ α2δ-1 interactions may control adult synapse formation and behavior, several of which we tested here.

First, we evaluated other possibilities that could explain why the loss of TSP1/2 leads to reduced performance. We found that TSP1/2-KO mice learn and adapt to changes in action-outcome contingencies when low effort is required for the reward. They also consumed similar amounts of food pellets and checked the food delivery port at the same rate as wild-type mice, so their food-seeking behavior was intact. We also ruled out hypoactivity because TSP1/2-KO mice presented no locomotion deficit during an open-field test to assess overall mouse movement in a new environment.

The loss of either TSP1/2 or α2δ-1 significantly reduces excitatory synapses in the developing brain [29,43]. This impairment has been previously shown to persist into adulthood for α2δ-1 KO mice [51,54]. Here, we show for the first time that adult TSP1/2-KO mice also have significantly reduced excitatory synapses in the ACC, as previously observed in α2δ-1 KO mice [54]. However, unlike in α2δ-1 KO mice, we observed a net increase in excitatory synapses in the ACC of TSP1/2-KO mice following instrumental training. This result strongly suggests that TSP1/2 are not the only synaptogenic α2δ-1 ligands required for training-induced synaptogenesis. Four isoforms of thrombospondins (TSP1 – TSP4) are expressed in the brain by astrocytes [43,45–47,94]. Therefore, TSP3 or TSP4 are likely signaling to α2δ-1 to promote training-induced synaptogenesis in the adult brain. Indeed, RNA-seq and qPCR analyses revealed a significant increase in Thbs4 expression in trained TSP1/2-KO mice which is highly likely to underlie the training induced synapse formation. Future studies investigating the functions of TSP3 and TSP4 in the adult CNS are needed to test this possibility.

Our analysis of gene expression changes shows that TSP4 is upregulated in the ACC of TSP1/2-KO mice following instrumental training, further suggesting that TSP4-α2δ-1 interaction could be causing training-induced synapse formation. Indeed, TSP4 is expressed by astrocytes in the adult brain and has been shown to promote excitatory synaptogenesis through its interaction with α2δ-1 [43,49].

An unexpected synaptic phenotype in the TSP1/2-KOs is the increase in inhibitory synapse numbers and function. In untrained TSP1/2 mice, genes associated with inhibition were highly upregulated compared to WT mice, a difference which was then downregulated in TSP1/2-KO ACC with operant training. This decrease in inhibition, coupled with the increase in excitation in the ACC of TSP1/2-KOs after training, suggests a major increase in overall excitation, which would explain why the TSP1/2-KO mice do not press the lever as much as the WT. We previously found that the increased excitation in the ACC, especially on neurons projecting to the dorsomedial striatum, decreases effort exertion and performance [54]. Therefore, an increased level of overall excitation in TSP1/2-KOs following training may explain the reduced performance by TSP1/2-KOs compared to WT.

Thrombospondins are expressed in several tissues and organs during development, including the heart, lung, skeletal muscle, brain, and bone [93,104,105]. Moreover, TSP1/2 is highly expressed in the developing cortex but not high in adults. Considering this, we wondered whether the behavioral effects can be specifically attributed to the loss of TSP1/2 in astrocytes. To specifically disrupt Thbs1 and Thbs2 genes in astrocytes, we used a viral CRISPR/Cas9 method. The ablation of TSP1/2 in adult astrocytes did not alter performance during operant tasks. However, when we ablated TSP1/2 in neonatal astrocytes, targeting a period corresponding to the start of astrocyte morphogenesis, synapse development, and normal expression of TSP1/2 in the cortex [106,107], we found a significant decrease in performance during high-effort tasks. This is consistent with previous findings in the literature showing that cortical astrocytes express TSP1/2 within the first two weeks of post-natal development, but the expression is diminished by P21 [29]. TSP1/2 likely only plays a critical but transient role in cortical synaptogenesis during early development [29]. Therefore, the loss of TSP1/2 during early development may have prevented the establishment of synaptic connections required for proper performance during high-effort instrumental tasks. Interestingly, neurodevelopmental disorders linked to synaptic dysfunction, such as autism spectrum disorders, have also been shown to be associated with a reduction in attributes of goal-directed behaviors [6–8,108,109]. Our findings reveal a link between astrocyte-secreted synaptogenic cues during early brain development and the control of instrumental actions in adulthood. This suggests that the developmental disruption of synaptogenesis due to astrocyte dysfunction may underlie some impairments in goal-directed behaviors.

## Materials and Methods

### Animals, Housing, and Genotyping

All mice were used in accordance with the Duke Division of Laboratory Animal Resources (DLAR) and Institutional Animal Care and Use Committee (IACUC) oversight (IACUC Protocol Numbers A173-14-07, A147-17-06, A263-16-12, A179-17-07, A117-20-05, A135-20-06, A110-23-04). The mice were kept under standard 12-hour day/night conditions. Wild type C57BL/6J (Stock #000664), B6.129S2-Thbs1tm1Hyn/J or TSP1-KO (Stock #006141), B6;129S4-Thbs2tm1Bst/J or TSP2-KO (Stock #006238), and B6J.129(B6N)-Gt (ROSA)26Sortm1(CAG-cas9*, EGFP)Fezh/J (JAX stock #026175) lines were obtained from Jackson labs. TSP1/2-KO mice were generated through a series of crossings between TSP1-KO and TSP2-KO mice. Male and female mice, between 3 and 5 months old, were acclimated to the researcher through 5 to 10 minutes of daily handling for one week. The animals were then food-restricted for 3-5 days until they reached 85-90% of their normal body weight, which was maintained by daily feeding with 1.5/2 grams of home chow after training.

### Instrumental Training

All instrumental training was performed as previously described [54]. The lever press training task was performed in operant chambers with light-resistant and sound-attenuating walls. Each chamber had a food magazine with an infrared beam, a food dispenser, two retractable levers, and a house light. The left lever was used for training, and a computer with a Med-PC-IV program controlled the setup and recorded animal behavior. Custom programs were used to analyze the collected data (available upon request).

### Fixed Ratio Schedule

To test the ability of mice to learn new behaviors, a Fixed Ratio 1 (FR1) continuous reinforcement schedule was used. Mice received one food pellet (Bio-Serv 14mg Dustless Precision Pellets) per lever press during a 3-day FR1 period, with sessions ending after 120 minutes or 50 rewards. This was followed by two additional testing schedules: a 3-day FR5 (one pellet for every 5 lever presses) and a 3-day FR10 (one pellet for every 10 lever presses), with sessions ending after 60 minutes or 50 rewards. We utilized Matlab to identify the start and end of lever press bouts based on the average inter-press interval probability distribution of the WT or control condition. After 9 days of testing, mice were sacrificed for histological analysis or used in the next behavioral test phase.

### Progressive Ratio, Extinction, and Omission Tests

We utilized a Progressive Ratio (PR) schedule to assess the level of effort mice are willing to expend to get a food reward. The PR schedule involved increasing the number of lever presses required for each subsequent reward earned. For our experiment, we increased the progression by 5. In this way, the first reward required 1 lever press, then the next reward required 6 lever presses, and this progressive increase in lever presses for reward delivery continued until the end of the session. The session ended after 60 minutes of the task; the lever was retracted, and the light in the Skinner box was turned off.

During extinction, the mice had access to the lever, but pressing the lever did not deliver a food reward. The session ended after 30 minutes; the lever was retracted, and the light in the Skinner box was turned off. During the omission schedule, the food reward was delivered every 20 seconds unless the mouse pressed the lever. Each time the mouse pressed the lever, the timer for reward delivery resets, which delays reward delivery.

### Open Field Test

After completing the operant task on food restriction, mice were provided with unlimited access to food and water for 2-3 days until they regained their normal body weight. The mice were then assessed in an open field chamber equipped with a blackfly camera (Flyr System, BFS-U3-04S2M-CS) to monitor their movements. The Bonsai software (https://bonsai-rx.org/) was utilized to automatically detect and record the x and y coordinates of the mouse’s center of mass. The testing session started after the mice had acclimated to the new room for 30 minutes. The mouse was then positioned in the center of the arena at the start of the recording, which was concluded after 30 minutes. The chamber was meticulously cleaned to prevent the influence of other animals’ scent stimuli, and the bedding was changed between different groups of littermates.

### Immunohistochemistry

Mice were anesthetized and perfused with Tris-Buffered Saline (TBS), followed by 4% Paraformaldehyde (PFA). Mouse brains were kept in 4% PFA overnight at 4°C, rinsed with TBS and stored in 30% sucrose in TBS for cryoprotection. The brains were then embedded in a 30% sucrose and Tissue Tek O.C.T. compound mixture and stored at −80°C. Finally, brains were cut into 25-30 µm coronal sections using a cryostat and stored in 50% glycerol in TBS. Brain sections were washed and permeabilized in TBS containing 0.2% Triton-X 100 (TBST). They were then blocked in TBST with 5% Normal Goat Serum for 1 hour at room temperature. Primary antibodies, including guinea pig anti-VGlut1 (1:2000 [AB5905, Millipore, MA]) and rabbit anti-PSD95 (1:300 [51–6900, Invitrogen, CA]), chicken anti-green fluorescent protein (1:1000 [GFP-1020, Aves Labs, CA]), Guinea pig anti-VGAT (Synaptic Systems #131004), and Mouse anti-gephyrin (Synaptic Systems #147-011), were diluted in TBST with 5% NGS, and sections were incubated overnight at 4°C. Subsequently, secondary Alexa-fluorophore-conjugated antibodies were added in TBST with 5% NGS for 2 hours at room temperature. Isotype-specific secondary antibodies were used for primary antibodies produced in mice to minimize background staining. The sections were washed in TBST and mounted on slides with Vectashield with DAPI (Vector Laboratories, CA).

To stain for Sox9, Olig2, NeuN, and DAPI, 25-30 µm coronal sections were washed and permeabilized in TBS containing 0.2% Triton-X 100 (TBST). They were then blocked in TBST with 5% Normal Goat Serum for 1 hour at room temperature. Primary antibodies, including rabbit anti-Sox9 (1:2000 [Millipore #AB5535]), Mouse anti-Olig2 (Clone 211F1.1; 1:500 [Millipore #MABN50]), and Mouse anti-NeuN (Clone A60; 1:1000 [Millipore #MAB377]) were diluted in TBST with 5% NGS, and sections were incubated overnight at 4°C. Subsequently, secondary Alexa-fluorophore-conjugated antibodies were added in TBST with 10% NGS for 3 hours at room temperature. DAPI (1:50,000) was added 15 minutes before the end of the 3-hour incubation. Isotype-specific secondary antibodies were used for primary antibodies produced in mice to minimize background staining. The sections were washed in TBST and mounted on slides with STED mounting media (20 mM Tris (pH 8.0), 0.5% N-propylgallate, 90% glycerol).

### Synapse Quantification

Synapse quantification was performed as previously described [54,64,110]. Briefly, synapse analysis was performed using immunohistochemistry on three independent brain sections per group. Confocal z-stacks of the ACC and DMS were imaged at 60× magnification and 1.64x zoom on an Olympus Fluoview 3000 confocal laser-scanning microscope. Genotype-blind analyses were conducted using an ImageJ macro called SynBot (https://github.com/Eroglu-Lab/Syn_Bot) to count colocalized synaptic puncta [65]. This quantification method accurately estimates the number of synapses by measuring co-localization rather than stained pre- or postsynaptic proteins.

The quantification method involves processing 1 µm thick maximum projections, subtracting backgrounds, and determining thresholds to detect discrete puncta without introducing noise. SynBot uses an algorithm to detect the number of puncta in close proximity across channels to quantify co-localized puncta.

### Whole-Cell Patch-Clamp Recording

For whole-cell patch-clamp recordings, 2-4 mice were used to miniature inhibitory postsynaptic current (mIPSC) for each genotype. During all recordings, brain slices were continuously perfused with standard aCSF at RT (∼25°C) and visualized by an upright microscope (BX61WI, Olympus) through a 40x water-immersion objective equipped with infrared-differential interference contrast optics in combination with a digital camera (ODA-IR2000WCTRL). Patch-clamp recordings were performed using an EPC 10 patch-clamp amplifier controlled by Patchmaster Software (HEKA). Data were acquired at a sampling rate of 50 kHz and low-pass filtered at 6 kHz.

To prepare acute brain slices, after decapitation, the brains were immersed in ice-cold artificial cerebrospinal fluid (aCSF, in mM): 125 NaCl, 2.5 KCl, 3 mM MgCl2, 0.1 mM CaCl2, 10 glucose, 25 NaHCO3, 1.25 NaHPO4, 0.4 L-ascorbic acid, and 2 Na-pyruvate, pH 7.3-7.4 (310 mOsmol). Coronal slices containing the ACC were obtained using a vibrating tissue slicer (Leica VT1200; Leica Biosystems). Slices were immediately transferred to standard aCSF (37°C, continuously bubbled with 95% O2 – 5% CO2) containing the same as the low-calcium aCSF but with 1 mM MgCl2 and 1-2 mM CaCl2. After 30 min incubation, slices were transferred to a recording chamber with the same extracellular buffer at room temperature (RT: ∼25°C).

To measure mIPSC, the internal solution contained the following (in mM): 77 K-gluconate, 77 KCl, 10 HEPES, 1 EGTA, 4.5 MgATP, 0.3 NaGTP, and 10 Na-phosphocreatine, pH adjusted to 7.2 – 7.4 with KOH and osmolality set to ∼ 300 mosM. mIPSCs were measured in the aCSF bath solution containing 1 µM tetrodotoxin and 10 µM 6-cyano-7-nitroquinoxaline-2,3-dione (CNQX), and 50 µM D-2-amino-5-phosphonopentanoate (D-AP5) at −70 mV in voltage-clamp mode. To measure mEPSC, the internal solution contained the following (in mM): 125 K-gluconate, 10 NaCl, 10 HEPES, 0.2 EGTA, 4.5 MgATP, 0.3 NaGTP, and 10 Na-phosphocreatine, pH adjusted to 7.2 – 7.4 with KOH and osmolality set to ∼ 300 mosM. mEPSCs were measured in the aCSF bath solution containing 1 µM tetrodotoxin and 50 µM Picrotoxin at −70 mV in voltage-clamp mode. Series resistance was monitored throughout all recordings, and only recordings that remained stable over the recording period (≤30 MΩ resistance and <20% change in resistance) were included. mIPSCs and mEPSCs were analyzed using Stimfit software (https://github.com/neurodroid/stimfit) [111]. To analyze the frequency, events were counted over 5 minutes of recording. At least 100 non-overlapping events were detected and averaged to obtain the average events for each cell. The peak amplitudes of the average mIPSC and mEPSC were measured relative to the baseline current, and only events larger than 5 pA were included. All chemicals were purchased from Sigma-Aldrich or Tocris.

### Cell Counting Analysis

The counting of DAPI, Sox9, Olig2, and NeuN-labeled cells was performed as previously described [35]. Tile scan images (11 z-stacks with 1µm step-size; 10µm total) containing the anterior cingulate cortex from adult WT and TSP1/2-KO mice were acquired rapidly at 30X objective using the resonant scanner of an Olympus FV 3000. The resolution of the stitched images was digitally restored to improve resolution using a trained content-aware image restoration (CARE) network [112]. The ROI of stitched images used for quantifying cell numbers spanned L1 through L6 of the ACC (568µm x 1072µm). A machine-learning-based method (Falk et al., 2019) was used to segment and identify cells labeled by each marker. The complete source code for this method is available at https://github.com/ErogluLab/CellCounts. 2 – 4 images per animal was used for segmentation and analysis. Segmented images were reviewed and adjusted, when necessary, for accuracy.

### Preparation of Samples for Sequencing

Tissue samples were harvested from untrained WT, untrained TSP1/2-KO, and trained TSP1/2-KO adult mice (8 animals per genotype, including 4 males and 4 females) within 60 minutes of the last behavior session or exposure to the operant chamber. Animals were anesthetized with 200 mg/kg tribromoethanol (avertin) and perfused with Tris-Buffered Saline (TBS, 25 mM Tris-base, 135 mM NaCl, 3 mM KCl, pH 7.6). The brain was immediately extracted, and the anterior cingulate cortex (ACC) was dissected before it was flash-frozen in liquid nitrogen and stored at −80°C until RNA extraction.

### RNA Extraction

Frozen samples were homogenized in 1000μl TRIzol Reagent (15596026, Thermo Fisher Scientific, Waltham, MA) and vortexed at 2000 rpm for 5 min. 200μl of chloroform (Sigma-Aldrich, C2432, St. Louis, MO) was added to each tube and vortexed for 2 min. Samples were allowed to phase separate before being centrifuged at 11,900 rpm for 15 min at 4°C, after which the top clear aqueous phase was separated into a fresh tube.

500μl of Isopropanol (Thermo Fisher Scientific, NY) was added, and samples were vortexed at 2000 rpm for 1 min, incubated at room temperature for an additional 10 mins, and then centrifuged for 10 mins. The supernatant was discarded, and the RNA pellet was washed two times with 1 ml of ice-cold 75% ethanol, air-dried, and resuspended in 40μl of RNase-free water. RNA samples were treated with DNaseI, cleaned, and concentrated using the Zymo RNA Clean & Concentrator Kit (Zymo Research R1013). A minimum of 1000ng of RNA was obtained from each tissue sample.

### Library Prep, Sequencing, and Data Analysis

All RNA samples were coded numerically. Sequencing was performed blind to sample identity by MedGenome. RNA-sequencing libraries were generated using the Illumina TruSeq stranded mRNA kit. The NovaSeq 6000 produced 40-72 million, 2×100 reads per replicate. Trimmomatic (v0.38) was used to trim the raw reads of adapters, and STAR (v1.6.3) [113] was used to align the reads to the reference mouse genome (mm10). Subread (featureCounts v1.6.3) was used to count the reads, and edgeR (v3.30.3) was used to conduct differential gene expression analysis in R. ShinyGo 0.80 [114] was used to identify GO terms for differentially expressed genes. RNA-sequencing data have been deposited in the Gene Expression Omnibus (GEO) repository with accession number: <u>GSE257869</u>.

### Single cell RNAseq comparison

We used the R package scMappR [88] to compare our RNA-sequencing data with single cell RNA-sequencing data. Significant genes above or below a fold change of 1.5 were used as input for the function *tissue_scMappR_internal*, using a cortical single cell dataset [87], accession number SRA667466.

### cDNA preparation and Real-time qPCR

RNA was extracted from untrained and trained TSP1/2-KO ACC tissue. cDNA libraries were generated by incubating the RNA samples in a solution containing qScript cDNA SuperMix (VWR #101414-102) and nuclease-free water. The incubation conditions were 25°C for 5 min, 42°C for 30 min, and 85°C for 5 min. The cDNA generated was diluted 10-fold with nuclease-free water to a final volume of 100µL per sample and stored at −80°C.

For real-time qPCR, cDNA samples were added to 96-well qPCR plates and incubated with Taqman™ Fast Advanced Master Mix (Applied Biosystems #4444556), nuclease-free water, and Taqman™ Gene Expression Assay (FAM) at a ratio of 5µL Master mix: 2.5µL water: 0.5µL Assay: 2µL sample. Three replicates per sample were used for qPCR analysis. A no-template control sample containing all reagents except cDNA was used as the negative control. Cycle threshold (Ct) values for the gene of interest were normalized to Gapdh as a housekeeping gene. The following Taqman™ Gene Expression Assays were used: Mouse Thbs4 (Mm03003598_s1, Applied Biosystems #4448892), Mouse Gapdh (Mm99999915_g1, Applied Biosystems #4331182).

### RNA fluorescent in situ hybridization (RNA-FISH)

Three to four 20µm sections containing the anterior cingulate cortex were directly mounted onto glass slides, kept at −80°C, and utilized for analysis less than one month after storage. HCR™ probes for Gad1 and Gad2 were purchased from Molecular Instruments. All steps were performed as described in the manufacturer’s protocol. Briefly, sections on slides were fixed in ice-cold 4% PFA for 15 min at 4°C, then immersed in a series of ethanol solutions (50%, 70%, and 100%) at room temperature for 5 min each, followed by a wash in PBS. A barrier was drawn around the tissue with a hydrophobic pen. The slides were dried, 200µL probe hybridization buffer was added to the tissues and incubated in a humidified chamber at 37°C for 10 min. The buffer was removed and replaced with probe solution, which was prepared by adding 0.4 pmol of each probe set to 100µL of probe hybridization buffer. The slides were incubated overnight at 37°C. To remove excess probes, the sections were incubated at varying proportions of probe wash buffer and 5x SSCT for 15 min, followed by a final incubation in 100% 5x SSCT for 15 minutes at 37°C and 5 min at room temperature. After drying the slides, 200µL of amplification buffer was added to the sections and incubated at room temperature for 30 minutes. The amplification buffer was removed and replaced with an amplification buffer containing the appropriate snap-cooled hairpins. The slides were incubated at room temperature overnight. Finally, the sections were washed three times in 5x SSCT for a total of 65 min, stained with DAPI in PBS for 3 minutes, and mounted with STED mounting media.

Tile scan images of the ACC were obtained at 60X magnification at 1µm step size for a total of 5µm z-stack on an Olympus FV 3000 microscope. Tile scan images were stitched using the Olympus imaging software, and images were max projected in ImageJ. Gad1+ and Gad2+ cells were manually annotated in the CellProfiler software [115]. DAPI was used to determine the total number of cells in the ROI and the percentage of cells labeled with *Gad1* or *Gad2*. The integrated intensity of Gad1 or Gad2 per cell was determined by summing all the pixel intensities within each Gad1+ or Gad2+ cell.

### Plasmids and CRISPR guides

To generate PGEARBOCS, the Cre expression cassette was cloned first into the pZac2.1-GfaABC1D-Lck-GCaMP6f (A gift from Dr. Baljit Khakh; Addgene plasmid #52924) by replacing Lck-GCaMP6f. Human U6 expression cassette and donor insertion sites (DIS) were synthesized as gBlocks (IDT) and cloned upstream of the gfaABC1D promoter. All the gRNAs used in this study were cloned into either the Sap1 site in the Human U6 expression cassette. To make the GEARBOCS donors, mCherry donors were PCR amplified, and HA donors were oligo annealed with overhangs to clone into the DIS. The Sanger sequencing protocol confirmed all generated plasmids. The sgRNA sequences used in this study were as follows; TSP1: 5’-AACTTGTCATCCGGCACAGCGG-3’, TSP2: 5’-GCCCGCCTACCGTTTTGTACGG-3’

### Production of Adeno-Associated Virus (AAV)

The AAV vectors AAV-PGEARBOCS-TSP1-KO and AAV-PGEARBOCS-TSP2-KO were generated and purified using a previously described method [116]. Briefly, HEK 293T cells were co-transfected with 15 µg of AAV vector, 30 µg of pAD-delta F6 helper plasmid, and 15 µg of capsid plasmids. Five 15 cm tissue culture dishes, each containing 12 × 10^6 HEK cells, were utilized for virus production. After three days post-transfection, cells were lysed using a buffer comprising 3 ml 5 M NaCl, 5 ml 1 M Tris-HCl (pH 8.5), and dH2O adjusted to pH 8.5 with NaOH. AAV particles were liberated through freeze/thaw cycles and treated with Benzonase at 37°C for 30 minutes. The purification process included iodixanol gradient ultracentrifugation, where cell lysates were loaded onto iodixanol gradients (15%, 25%, 40%, and 60%) in OptiSeal tubes and centrifuged at 67,000 rpm for 1 hour at 18°C. AAV particles were harvested from the 40%-60% iodixanol interface, mixed with ice-cold DPBS, and concentrated using a Vivaspin column (100 MWCO) at 4°C.

### Retro-orbital and Intracerebroventricular AAV injections

For retro-orbital injections, adult B6J.129(B6N)-Gt (ROSA)26Sortm1(CAG-cas9*, EGFP)Fezh/J (Cas9-EGFP) mice were placed in a stereotaxic frame, and anesthetized through inhalation of 1.5% isoflurane gas. 10μl of purified AAVs (titer of ∼1 × 10^12^ GC/ml) was intravenously injected into the retro-orbital sinus. The intracerebroventricular injection was performed as previously described [117]. Following hypothermic anesthesia, the respective AAV vectors were administered to Cas9-EGFP mouse pups on postnatal day one (P1). The injection site is approximately 0.8-1 mm lateral from the sagittal suture, halfway between lambda and bregma. 1.5μl of purified AAVs (titer of ∼1 × 10^12^ GC/ml) was injected per hemisphere using a 32G needle (Hamilton, Reno, NV, USA).

### Quantification and Statistical Analysis

Statistical analyses were performed using GraphPad Prism 8, 9, and 10 software. The sample size and specific statistical tests for each experiment can be found in the corresponding figure legends. Exact adjusted p-values are provided in the figures for each experiment; when not present, the difference was not significant (p>0.05). Detailed information on the statistical analysis is included in the figure legends. Significance levels for all quantifications are as follows: *p<0.05, **p<0.01, ***p<0.001, and ****p<0.0001. Sample sizes were determined based on prior experience with each experiment to ensure a high power for detecting specific effects. No predetermined statistical methods were used to establish the sample size.

## Acknowledgments

This work was supported by grants from the National Institutes of Health (NS096352 and AG059409 to C.E., DA040701, NS094754 to H.H.Y., NS112565 to K.S.).

F.P.U.S. was supported by postdoctoral fellowships from the Regeneration Next Initiative and the Ramon y Cajal Young Investigator Award (RYC2021-033202-I). S.W. was supported by postdoctoral fellowships from the Foerster-Bernstein Family. Illustrations were made with BioRender.com. Thanks to Dr. William Wetsel for his critical feedback on the manuscript. C.E. is an HHMI Investigator.

## Author Contributions

Conceptualization, OOL and CE; Methodology, OOL, FPUS, SW, DSB, KS, SJ, HHY, CE; Investigation, OOL, FPUS, SW, DSB, KS, SJ; Formal analysis, OOL, FPUS, SW, DSB, KS; Resources, FPUS, FPUS; Writing – original draft, OOL and CE; Writing – Review & Editing, OOL, FPUS, SW, DSB, KS, SJ, HHY, CE, Funding Acquisition, HHY, CE.

---

**S1 Fig.**
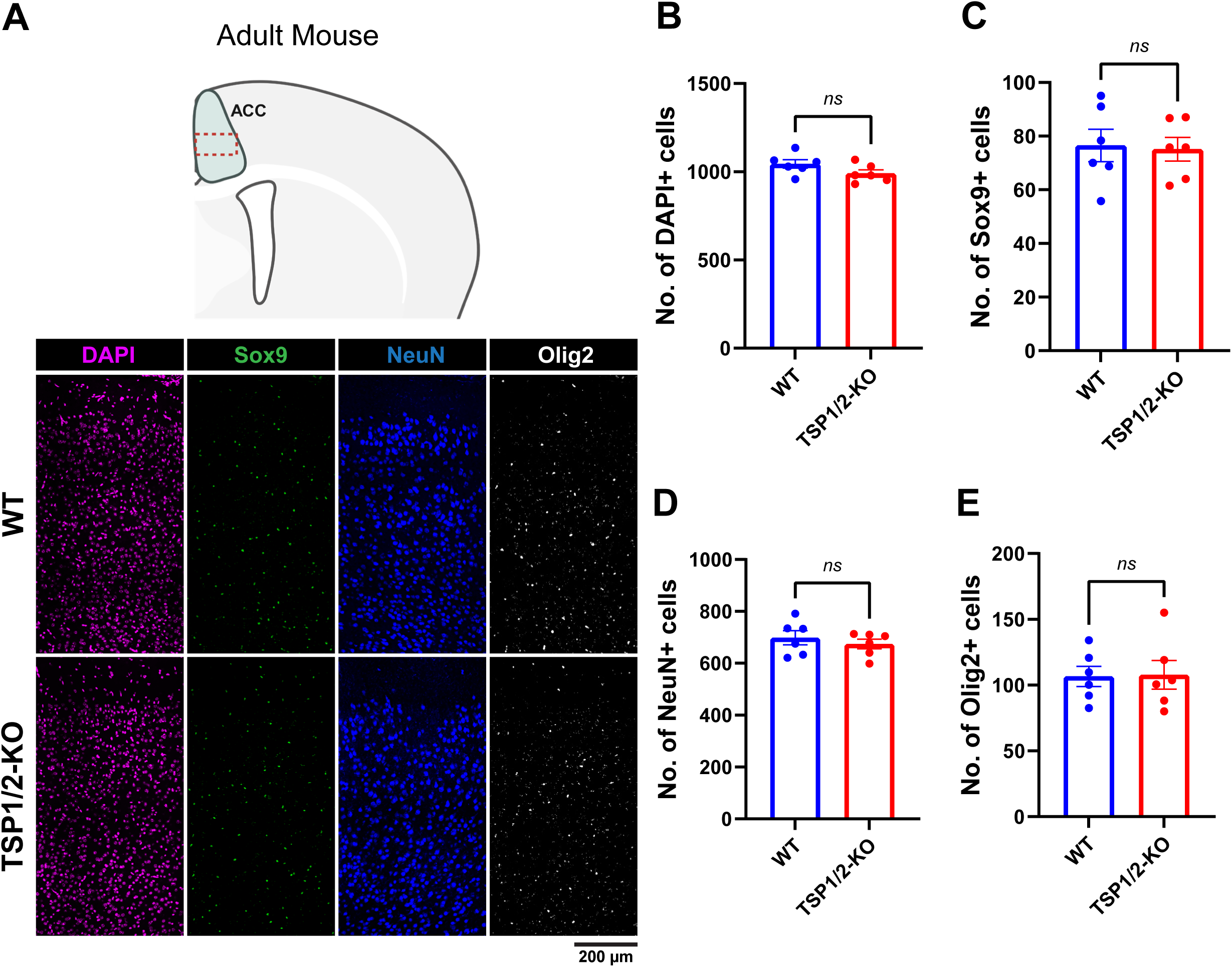
Loss of TSP1/2 does not alter neuronal and glial cell numbers in the ACC. **(A)** Top: Schematic representation of the ACC and the region of interest (red dashed rectangle) imaged for quantification. Bottom: Representative images showing the staining for cell markers DAPI, Sox9, NeuN, and Olig2 in the ACC of WT and TSP1/2-KO mice. **(B)** Quantification of the number of DAPI+ cells in the ACC of WT (n = 6; 3 male and 3 female; 1044 ± 23.83) and TSP1/2-cKO (n = 6; 3 male and 3 female; 990.3 ± 20.57) mice. Unpaired two-tailed T-test [t (9.792) = 1.707, p=0.1192]. **(C)** Quantification of the number of Sox9+ cells in the ACC of WT (n = 6; 76.51 ± 6.016) and TSP1/2-KO (n = 6; 75.13 ± 4.414) mice. Unpaired two-tailed T-test [t (9.174) = 0.1839, p=0.8581]. **(D)** Quantification of the number of NeuN+ cells in the ACC of WT (n = 6; 698.1 ± 26.95) and TSP1/2-KO (n = 6; 674 ± 18.74) mice. Unpaired two-tailed T-test [t (8.919) = 0.7336, p=0.4820]. **(E)** Quantification of the number of Olig2+ cells in the ACC of WT (n = 6; 106.6 ± 7.728) and TSP1/2-KO (n = 6; 107.8 ± 10.92) mice. Unpaired two-tailed T-test [t (9.006) = 0.0897, p=0.9305]. Data shown as mean ± s.e.m.

**S2 Fig.**
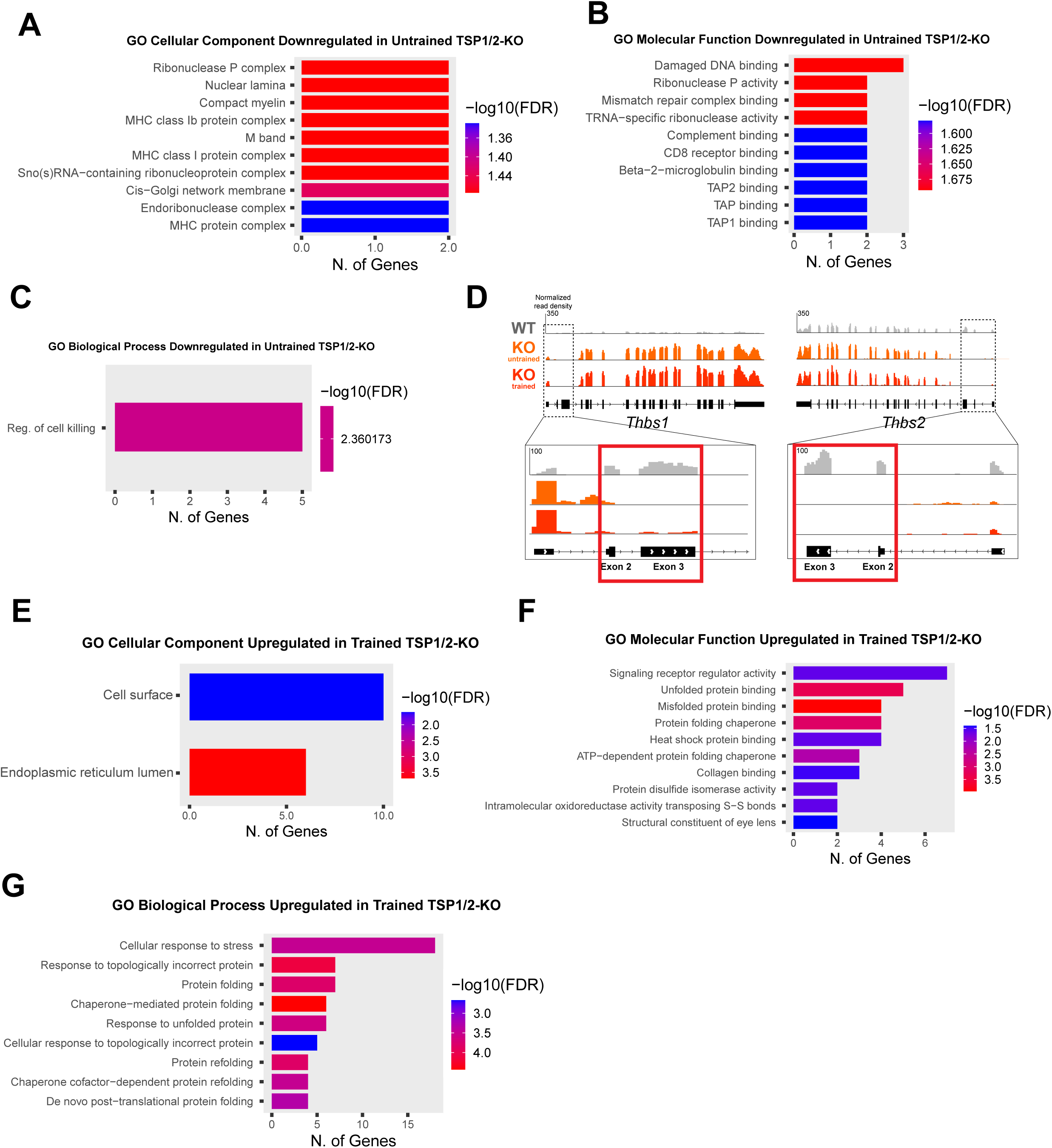
Gene expression changes in the ACC due to loss of TSP1/2 and instrumental training. **(A)** Gene Ontology (GO) Cellular Components for genes that are downregulated in untrained TSP1/2-KO compared to untrained WT (ShinyGo 0.80, FDR < 0.05). **(B)** Gene Ontology (GO) Molecular Function for genes that are downregulated in untrained TSP1/2-KO compared to untrained WT (ShinyGo 0.80, FDR < 0.05). **(C)** Gene Ontology (GO) Biological Process for genes that are downregulated in untrained TSP1/2-KO compared to untrained WT (ShinyGo 0.80, FDR < 0.05). **(D)** Example of exon level mRNA transcript abundance for Thbs1 and Thbs2 in WT, untrained and trained TSP1/2-KO RNA sequencing data. **(E)** Gene Ontology (GO) Cellular Components for genes that are upregulated in trained TSP1/2-KO compared to untrained TSP1/2-KO (ShinyGo 0.80, FDR < 0.05). **(F)** Gene Ontology (GO) Molecular Function for genes that are upregulated in trained TSP1/2-KO compared to untrained TSP1/2-KO (ShinyGo 0.80, FDR < 0.05). **(G)** Gene Ontology (GO) Biological Process for genes that are upregulated in trained TSP1/2-KO compared to untrained TSP1/2-KO (ShinyGo 0.80, FDR < 0.05).

**S3 Fig.**
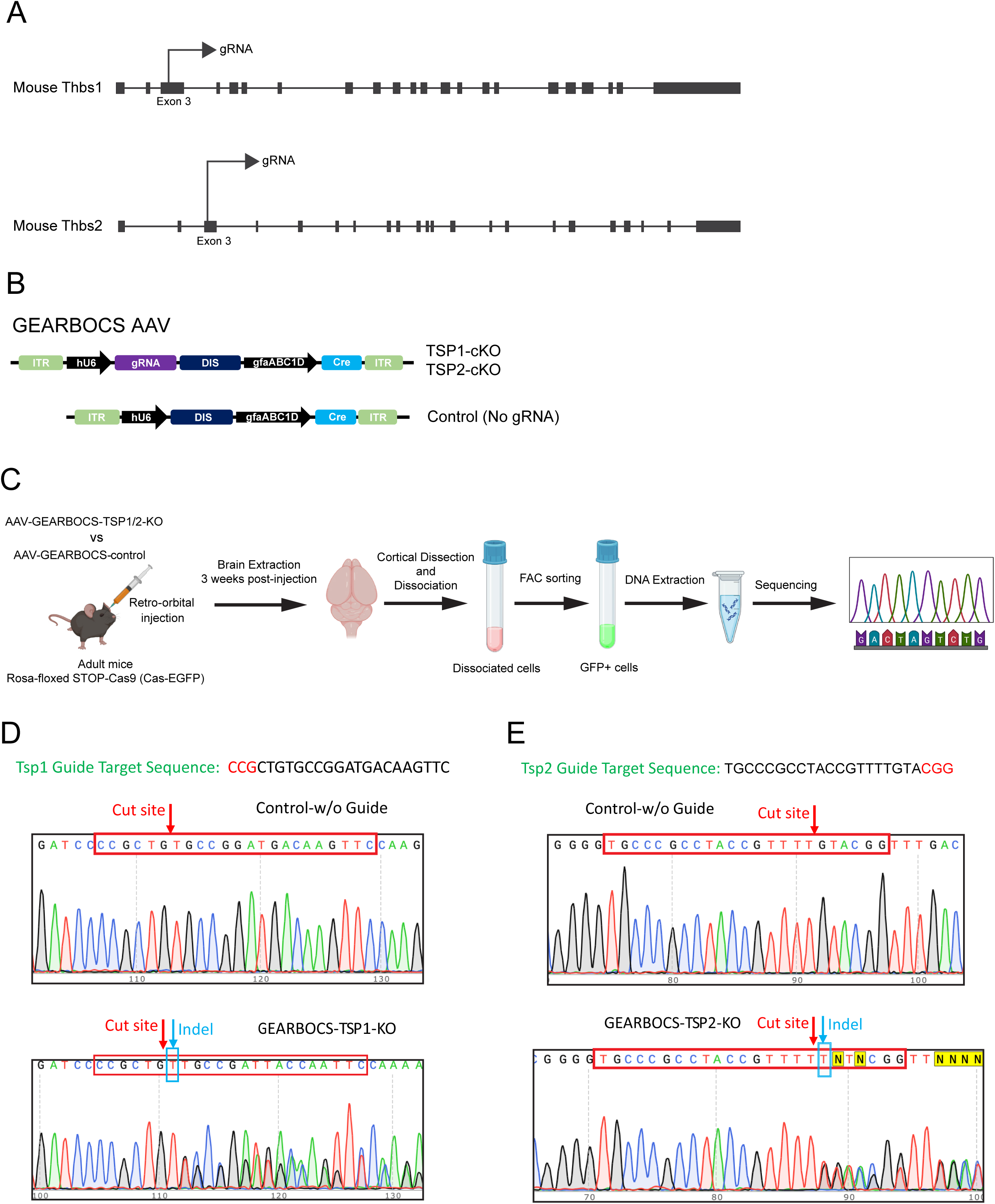
Strategy for specific ablation of thrombospondins 1 and 2 in astrocytes. **(A)** Schematic representation of the guide RNA target sites within the Thbs1 and Thbs2 locus, which codes for TSP1 and TSP2, respectively. **(B)** Schematic of AAV-GEARBOCS-TSP1-KO, AAV-GGEARBOCS-TSP2-KO, and the control plasmid, which does not contain any guide RNA. **(C)** Schematic representation of the protocol for isolating AAV transduced cells from retro-orbitally injected mice and the extraction of DNA for sequencing. **(D)** Top: A chromatogram showing the DNA sequence at the TSP1 targeted locus of GFP+ cells from mice injected with the control vector. Bottom: A chromatogram showing the insertion of indels into the same locus of GFP+ cells from mice injected with TSP1 knockout viral vector. **(E)** Top: A chromatogram showing the DNA sequence at the TSP2 targeted locus of GFP+ cells from mice injected with the control vector. Bottom: A chromatogram showing the insertion of indels into the same locus of GFP+ cells from mice injected with TSP2 knockout viral vector.

**S4 Fig.**
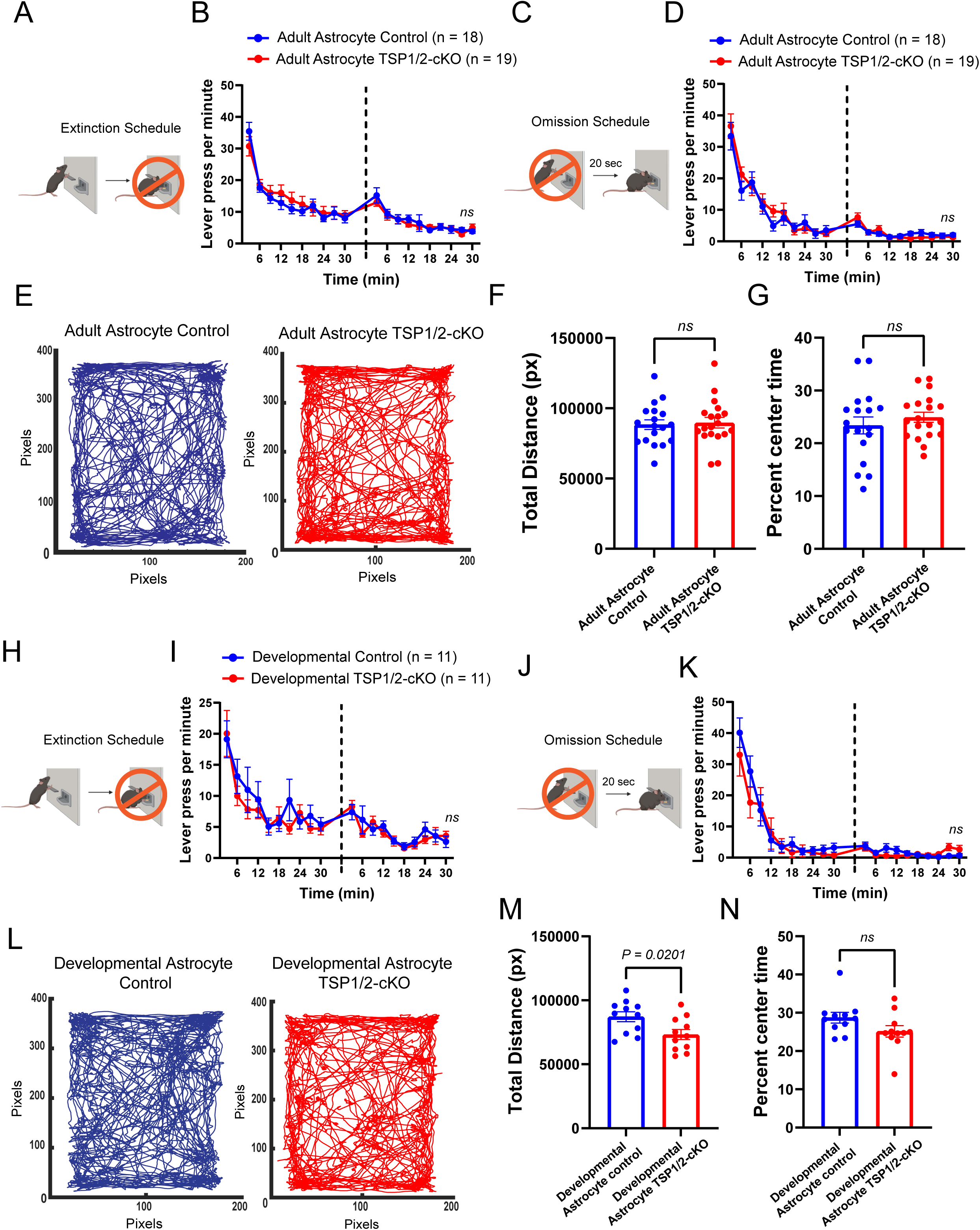
Loss of TSP1/2 in neonatal or adult astrocytes does not impact extinction or omission in adult mice. **(A)** Schematic representation of the extinction schedule. **(B)** Lever press per minute in 3 min bins for the 2 days of testing (dashed line) during extinction schedule for adult control (n =18) and TSP1/2-cKO (n = 19) animals. Repeated measures Two-way ANOVA. No main effects of genotype [F (1, 35) = 0.017, p = 0.897], main effect of Time [F (8.680, 303.8) = 37.77, p < 0.0001], and no effect of interaction [F (19, 665) = 0.742, p = 0.777]. **(C)** Schematic representation of the omission schedule. **(D)** Lever press per minute in 3 min bins for the 2 days of testing (dashed line) during omission schedule (n =18) and TSP1/2-cKO (n = 19) animals. Repeated measures Two-way ANOVA. No main effects of genotype [F (1, 35) = 0.187, p = 0.668], main effect of Time [F (5.619, 196.7) = 50.14, p < 0.0001], and no interaction [F (19, 665) = 0.744, p = 0.774]. **(E)** Example of open field patterns of locomotion by adult control and TSP1/2-cKO mice. **(F)** Bar graph of the total distance traveled in pixels for adult control (n = 18; 8.84*104 ± 3.4*103 pixels) and TSP1/2-cKO (n = 19; 8.97*104 ± 3.8*103 pixels) mice during an open field test. Unpaired two-tailed T-test [t (35) = 0.263, p=0.7939]. **(G)** Percent of time spent in the center of the arena by adult control (n = 18; 23.39 ± 1.59%) and TSP1/2-cKO (n = 19; 24.93 ± 0.95%) mice during an open field test. Unpaired two-tailed T-test [t (35) = 0.838, p=0.4077]. **(H)** Schematic representation of the extinction schedule. **(I)** Lever press per minute in 3 min bins for the 2 days of testing (dashed line) during extinction schedule for developmental control (n =11) and TSP1/2-cKO (n = 11) mice. Repeated measures Two-way ANOVA. No main effects of genotype [F (1, 20) = 0.6275, p = 0.4376], main effect of Time [F (5.271, 105.4) = 13.91, p < 0.0001], and no interaction [F (19, 380) = 0.6151, p = 0.8953]. **(J)** Schematic representation of the omission schedule. **(K)** Lever press per minute in 3 min bins for the 2 days of testing (dashed line) during omission schedule for developmental control (n = 11) and TSP1/2-cKO (n = 11) mice. Repeated measures Two-way ANOVA. No main effect of genotype [F (1, 20) = 0.6054, p = 0.4456], main effect of Time [F (3.307, 66.13) = 29.96, p < 0.0001], and no interaction [F (19, 380) = 0.8390, p = 0.6598]. **(L)** Example of open field patterns of locomotion by developmental control and TSP1/2-cKO mice. **(M)** Bar graph of the total distance traveled in pixels for developmental control (n = 11; 8.72*104 ± 3.9*103 pixels) and TSP1/2-cKO (n = 11; 7.32*104 ± 3.9*103 pixels) mice during an open field test. Unpaired two-tailed T-test [t (20) = 2.526, p=0.0201]. **(N)** Percent of time spent in the center of the arena by control (n = 11; 28.71 ± 1.38%) and TSP1/2-cKO (n = 11; 25.15 ± 1.51%) mice during an open field test. Unpaired two-tailed T-test [t (19.86) = 1.742, p=0.0969]. Data shown as mean ± s.e.m.

**S1 File. List of GO terms for differentially expressed genes in the untrained and trained TSP1/2-KO ACC.**

**S2 File. Exon level differential expression for Thbs1 and Thbs2.**

**S3 File. Cell-type specificity of differentially expressed genes in trained compared to untrained TSP1/2-KO ACC.**

